# The gene regulatory effects of selective glucocorticoid receptor ligands

**DOI:** 10.1101/2025.06.16.659962

**Authors:** Nicholas Giroux, Graham Johnson, Alejandro Barrera, Timothy Reddy

**Affiliations:** Department of Biomedical Engineering, Pratt School of Engineering, Duke University, Durham, NC, 27708, USA; Duke Human Vaccine Institute, School of Medicine, Duke University Medical Center, Durham, NC, 27710, USA; Department of Biostatistics and Bioinformatics, School of Medicine, Duke University Medical Center, Durham, NC, 27710, USA; Center for Advanced Genomic Technologies, Duke University, Durham, NC, 27708, USA

## Abstract

Synthetic glucocorticoids (GCs), which induce the transcription factor activity of the glucocorticoid receptor (GR), are frequently prescribed anti-inflammatory therapeutics that have been in use for over 70 years. Despite their broad immunosuppressive utility, sustained use of GCs is often intolerable due to the prevalence of adverse side effects. A longstanding goal has been to make synthetic GCs safer by developing selective GR ligands that have similar anti-inflammatory activity but without the burden of side effects. To evaluate the ability of synthetic GCs to target specific subsets of the GC response, we completed a genome-wide comparative analysis of changes in gene expression and gene regulatory element activity in response to ten ligands with various evidence of dissociated adverse side effects. We measured the gene expression response using mRNA-seq and the gene regulatory element response using genome-wide STARR-seq. Effects associated with each ligand were highly correlated with and linearly related to the response to dexamethasone, a strong, non-selective GR agonist used as a positive control for this study. Furthermore, 93% of the variation in regulatory element activity responses could be explained by the efficacy of each ligand alone. We also found limited evidence of differential enrichment of chromatin context-specific markers of regulatory activity with each ligand. Based on those findings, we developed a simulation framework to evaluate selectivity of GR ligands. We conclude that the ligands we tested elicit attenuated molecular responses according to their respective efficacies, and do not selectively target subsets of the molecular GC response.

## Introduction

The ligand-inducible activity of the glucocorticoid receptor (GR), a member of the nuclear receptor family of transcription factors (TFs), regulates expression of thousands of genes - as much as 20% of the genome - implicated in development, metabolism, and immunity.^1,2^ GR responds to external stimuli, exerting pleiotropic effects through its interactions with the genome directly and secondary effects via regulation of other TFs including activator protein 1 (AP-1), GATA-binding factor 1 (GATA1), and nuclear factor-kappa B (NFκB).^3,4^ In response to inflammation, glucocorticoids (GCs), such as the endogenous human steroid hormone cortisol, coordinate cellular and molecular processes necessary for potent immunosuppression.^5^ To that end, synthetic GC therapies have been developed for a range of clinical applications, including use as treatments for autoimmune, rheumatic, and endocrine disorders as well as in chemotherapy or organ transplantation regimens.^6–9^ Indeed, nuclear receptors are the target of nearly 14% of all FDA-approved drugs.^10^ GC therapy use is highly prevalent, with studies estimating that one in five adult Americans received prescriptions for short-term courses of GCs.^11^

Systemic adverse side effects are common as a result of chronic or excessive exposure to GCs at pharmacological concentrations.^12,13^ Severe side effects from over-exposure to GCs include insulin resistance and hyperglycemia, hypertension, osteoporosis, impairment of normal wound healing, increased risk of infection, and glaucoma, due to increased intraocular pressure.^14–16^ The risk and severity of these effects increases with both duration of GC use and dosage. Those side effects severely limit the utility of GC therapies for many patients. Therefore, the development of safer and more effective compounds is an opportunity to substantially benefit millions of patients worldwide.

GR activity depends on the utilization of several functional surfaces, including the DNA-binding domain (DBD), ligand-binding domain, and two activation function (AF1 and AF2) domains, that cooperate to recruit coregulators and control the expression of target genes.^1,17^ Specifically, coregulator recruitment is determined by the conformation of GR; conformation changes, in turn, are induced by a number of allosteric effectors such as ligands and differences in DNA binding sequences.^18–23^ Together these signals produce specific patterns of GR activity that depend on cellular and genomic context. Indeed, patterns of gene-specific activation have been mapped to each functional domain by genetic studies that perturbed the conformation of GR.^24^ By disrupting the DBD with a single point mutation, for example, it was once proposed that dimerization is dispensable for the trans-repressive mode of GR activity.^25^ Later experiments improved that model by showing how dimerization and DNA binding stabilize interactions with GR coregulators and other TFs.^26–30^ For instance, two subunits of the Mediator complex, MED1/DRIP205 and MED14/DRIP150, have been shown to preferentially interact with AF2 in a ligand-dependent manner and AF1 independent of ligand binding, respectively, with different consequences for downstream gene regulation.^31–33^ Together, these findings show that there are many binding interfaces which could be selectively manipulated in efforts to tune GC responses.

Pharmacological efforts to develop selective GR modulators (SGRMs), theoretically capable of dissociating the therapeutic anti-inflammatory benefits of classical GC therapy from the adverse side effects, have been the focus of hundreds of scientific and clinical studies over several decades.^34^ Despite a growing number of candidate SGRM compounds, few have progressed beyond preclinical studies, and those tested in controlled clinical studies largely lack evidence of a reduction in adverse, often metabolic-related, effects or an equal or greater immunosuppressive efficacy.^30,35–37^ For example, just one of 14 investigational candidate SGRMs developed for arthritis was studied in a clinical trial setting and had a better balance between clinical efficacy and safety.^38^

Here, we tested the hypothesis that selective GR ligands elicit selective gene expression responses by activating and repressing specific subsets of gene regulatory elements. To do so, we evaluated two key molecular features using genome-wide approaches: changes in gene expression using mRNA-seq and changes in gene regulatory element activity using STARR-seq. We identified thousands of GR ligand-responsive genes and tens of thousands of GR ligand-responsive regulatory elements. Using those data, we estimated transcriptional and gene regulatory element efficacies for each compound and found that nearly all the variation in responses across ligands is well-explained by those efficacies. We developed a new simulation model based on those findings that can be to guide the development of more selective GC therapies. Taken together, these findings present an experimental and statistical framework for evaluating the selectivity of a wide range of drug responses that act via transcriptional regulation.

## Results

### Measurement of gene expression and regulatory element activity responses to classical GCs and SGRMs

To identify distinct patterns of GR activity induced by classical GCs and SGRMs, we used mRNA-seq to measure changes in gene expression and whole-genome STARR-seq to measure changes in regulatory element activity (Fig. 1A). We measured the genomic responses to 10 GR ligands in A549 human lung epithelial cells. We selected the ligands to include both GR agonists and antagonists as well as SGRMs that have evidence of dissociated trans-activation. Specifically, we selected two commonly used agonists as positive controls (dexamethasone and hydrocortisone), one antagonist (RU486), and seven SGRMs (agonists: AZD2906, AZD9567, GW870086, mapracorat, ZK216348; antagonist: CORT108297; modulator: Compound A). All compounds were dissolved in DMSO, and we used equal volume DMSO as a vehicle control. Properties of all ligands are detailed in Table 1. For all ligands, we measured the genomic responses after treatment at 1 μM for 4 hr.

**Figure 1.**
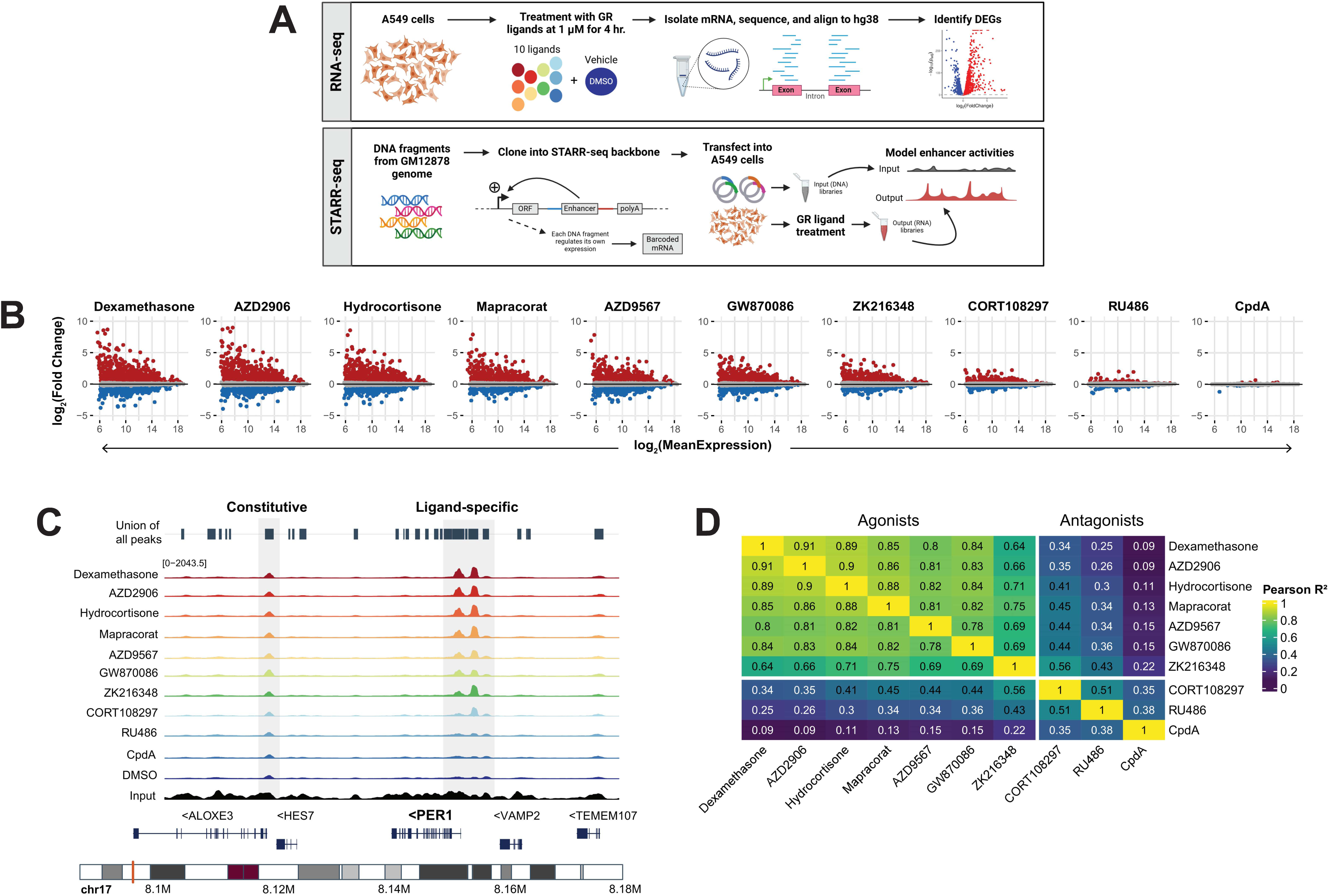
Genome-wide transcriptional and gene regulatory element activity effects in response to GR ligand treatment. A. Experimental design for the assay of gene expression and regulatory element responses to 10 classical and selective GR ligands. Created in BioRender. B. Distribution of significant differentially expressed genes in each ligand treatment condition. C. Ligand-specific and constitutive regulatory element activity at the *PER1* locus. D. Correlation between regulatory element activities at all 49,711 ligand-responsive STARR-seq regions.

**Table 1.** Properties of tested GR ligands.

| Compound | Activity | Selectivity | IC<sub>50</sub> (nM) | Clinical trial |
| --- | --- | --- | --- | --- |
| Dexamethasone | Agonist | No | 2.0 | Yes, phase 4 completed |
| AZD2906 | Agonist | Yes | 2.5 | No, pre-clinical |
| Hydrocortisone | Agonist | No | 15.3 | Yes, phase 4 completed |
| Mapracorat | Agonist | Yes | 3.9 | Yes, phase 3 completed |
| AZD9567 | Agonist | Yes | 3.8 | Yes, phase 2 completed |
| GW870086 | Agonist | Yes | 10.1 | Yes, phase 2 completed |
| ZK216348 | Agonist | Yes | 20.3 | No |
| CORT108297 | Antagonist | Yes | 0.9 | Yes, phase 2 completed |
| RU486 | Antagonist | Yes | 2.6 | Yes, phase 4 completed |
| Compound A (CpdA) | Modulator | Yes | 6.4 | No |
| DMSO | No activity | N/A | N/A | N/A |

The activity, selectivity, half-maximal inhibitory concentration (IC<sub>50</sub>), and status of clinical trial studies for each GR ligand were determined by reviewing relevant literature. Ligand activity was categorized as being either an agonist (N = 7), antagonist (N = 2), or modulator (N = 1). When multiple IC<sub>50</sub> measurements were available for any given ligand, we reported the average. When we identified multiple clinical trial studies for a ligand, we reported the most advanced progress available from ClinicalTrials.gov.

We hypothesized that we would observe ligand-specific changes in gene expression and regulatory element activity. Specifically, because SGRMs were designed in part to impair GR homodimer formation, it has been hypothesized that they would induce GR binding at a subset of response elements, thus leading to selective activation of regulatory elements, cofactor recruitment, and downstream target gene expression.^39^

### GR ligand-specific differential gene expression

To first establish which genes had GR ligand-inducible differential expression, we used mRNA-seq to compare gene expression levels between each treatment condition and the DMSO control. In total, we assayed four replicates for each treatment condition for a total of 44 mRNA-seq datasets. Treatment with the ligands we tested resulted in significant differential expression for 8,072 genes at a 5% false discovery rate (Figure 1B). Dexamethasone regulated the greatest number of those genes, with 4,818 (59.7%) having significant differential expression following treatment. Of those dexamethasone-responsive genes, 85% (N = 4,099) had a significant change in expression in at least one additional ligand treatment condition.

There was substantial consistency in the gene expression effects across ligands. Of the 8,072 differentially expressed genes, 3,883 (48.1%) had increased expression in all conditions where the effect was significant, 3,934 (48.7%) had decreased expression in all conditions where the effect was significant, and the remaining 255 (3.2%) had increased expression in some conditions and decreased expression in others. Additionally, of the genes responsive to each individual ligand treatment, the proportion that were also dexamethasone-regulated ranged from a minimum of 52.4% for Compound A to a maximum of 87.5% for ZK216348 (mean = 71.5% ± 11.5%).

Overall, AZD2906 and dexamethasone produced the strongest changes in gene expression, Compound A had little to no effect on gene expression, and the remaining ligands had intermediate effects. For example, the canonical glucocorticoid-activated gene *PER1* was upregulated 15-fold by AZD2906, 13-fold by dexamethasone, 6-fold by GW870086, and had no change in expression after Compound A treatment. Similarly, the AP-1 component *FOSL1* was repressed by 1.6-fold by AZD2906 and dexamethasone, 1.3-fold by GW870086, and also had no change in expression after Compound A treatment. Together, these results demonstrate that the SGRMs we tested can and do target dexamethasone-responsive genes in a consistent manner, albeit with different effect sizes. Combined with our finding that 96.8% of differentially expressed genes had effects in the same direction as dexamethasone, these patterns suggest that the ligands can activate and repress gene expression with similar specificity due, potentially, to shared regulatory mechanisms.

### GR ligand-specific differential regulatory element activity

GR regulates gene expression by modulating the activity of nearby gene regulatory elements.^40–42^ Reporter gene assays have been instrumental in revealing those changes in regulatory element activity, and previous studies have identified thousands of glucocorticoid-responsive gene regulatory elements.^39,43^ To measure ligand-specific changes in regulatory element activity genome-wide, we used STARR-seq, a high-throughput reporter assay (Fig. 1A, Fig. S1A).^44^ Specifically, we constructed a whole-genome STARR-seq assay library using DNA fragments from the GM12878 genome. The reporter library consisted of approximately 870 million unique fragments with an average length close to 1 kb (minimum: 0.75 kb; average: 0.96 kb; maximum: 2.50 kb). The 1 kb fragment length allows us to study larger and more complex regulatory elements than is typical for high-throughput reporter assay studies. Overall, the assay library had an average genome-wide coverage of 530 fragments per base, giving us very high resolution into the gene regulatory element responses to various GR ligands (Fig. S1B-C). For each ligand treatment, we assayed the STARR-seq library in four technical replicates for each of four biological replicates, generating a total of 176 STARR-seq datasets.

STARR-seq reveals both GR ligand-responsive regulatory activity and constitutive regulatory element activity. For example, there are two well-studied glucocorticoid-response elements flanking the *PER1* promoter.^45^ Both elements have significant glucocorticoid responsive activity in our STARR-seq assays, and the magnitude of the effect varies across ligands in a similar pattern as *PER1* gene expression (Fig. 1C). In contrast, there is a constitutively active gene regulatory element near the *ALOXE3* promoter ∼30 kb downstream of *PER1*. For the remainder of this study, we focus on the GR ligand-responsive regulatory elements, which we define as having a significant change in regulatory activity compared to the DMSO control.

We first called regulatory elements de-novo in each treatment condition by comparing the activity in that condition to the input library using CRADLE.^46^ In total, we identified 182,080 genomic regions with regulatory activity in A549 cells in any condition or in vehicle control at a 5% false discovery rate. We then tested those regions for evidence of ligand-specific regulatory activity. That yielded a union set of 49,711 (27%) genomic regions with significant changes in regulatory activity after any one of the GR ligand treatments assayed. We detected the largest number of ligand-specific regulatory elements with activity after dexamethasone treatment (N = 31,294). There was also substantial overlap in GR ligand-responsive regulatory activity between ligands. For example, of the dexamethasone-responsive regulatory elements, 85% (N = 26,541) had a significant change in activity in at least one additional ligand treatment condition.

Ligand-specific regulatory activity was highly consistent across conditions. Of the 49,711 elements with differential activity, 28,646 (57.6%) had increased activity in all conditions where the effect was significant, and 20,984 (42.2%) had decreased activity in all conditions where the effect was significant. Only 81 genomic regions (0.2%) had increased activity in some conditions and decreased activity in others. That fraction is well below our false discovery rate threshold, and we conclude those 81 regions are likely to be false positives in our statistical analysis.

The strong agonists dexamethasone and AZD2906 produced the strongest changes in regulatory element activities while Compound A had the weakest effects. The remaining ligands had intermediate effects. To demonstrate this trend, we identified three elements near the *PER1* locus with significant ligand-specific effects (Fig. S1D). Two of these elements were proximal to the *PER1* promoter, and the third was in the 5’ untranslated region (UTR). The activity of the 5’ UTR element was increased 3.6-fold by dexamethasone, 3.5-fold by AZD2906, 2-fold by GW870086, and had no significant change after Compound A treatment. Similarly, we identified a *FOSL1* promoter-proximal element with activity decreased 2-fold by dexamethasone, 1.9-fold by AZD2906, 1.7-fold by GW870086, and having no significant change after Compound A treatment (Fig. S1E).

A substantial fraction of the genomic regions with STARR-seq activity also had accessible chromatin, a common indicator of regulatory element activity.^47^ Specifically, we identified 98,869 and 111,123 sites of chromatin accessibility before and after dexamethasone treatment, respectively. Nearly all those chromatin accessible sites were the same between conditions (N = 76,262). That overlap is consistent with previous studies demonstrating that most GR binding activity targets regions with chromatin accessibility prior to ligand treatment.^48,49^ Of those sites with chromatin accessibility after dexamethasone treatment, 37,067 (34%) also had regulatory activity in STARR-seq in the same conditions. Conversely, 20% of the genomic regions identified with STARR-seq also had chromatin accessibility. We interpret those sites as likely regulatory elements in A549 cells, whereas we interpret the sites with regulatory activity in STARR-seq but undetected chromatin accessibility as potential latent regulatory elements active in other cell types.

GR ligand-responsive regulatory activity was highly correlated across the 49,711 ligand-responsive STARR-seq regions. The average correlation between these regulatory element activities was R^2^ = 0.62 ± 0.31 (Fig. 1D). Moreover, regulatory activity was especially well correlated among the subset of six GR agonists (AZD2906, hydrocortisone, mapracorat, AZD9567, GW870086, and ZK216348) and dexamethasone (average R^2^ of 0.82 ± 0.10). Those correlations were significantly higher than for the antagonists/modulators (CORT108297, RU386, and CpdA; average R^2^ of 0.31 ± 0.13; p = 0.005, Welch’s t-test). We confirmed that the correlations calculated for the six agonistic ligands clustered together, and we interpret these results to indicate that, on average, most variation in regulatory element activity (82%) can be explained relative to the effect that the strong agonist dexamethasone has on those elements (Fig. S1F).

### Modulation of transcription by GR ligands compared to dexamethasone

To evaluate the effects of the various ligands on gene expression, we estimated differential gene expression from the previously described RNA-seq assays. Both the magnitude of the responses and the number of statistically significant differentially expressed genes (DEGs) varied widely (Fig. 2A). Similar to what we observed with STARR-seq, we detected the most DEGs after treatment with dexamethasone (4,818) and AZD2906 (5,240), and we detected the fewest DEGs in response to Compound A (21). Across all ligands, we observed an average of 2,980 DEGs and a median of 3,606 DEGs. We refer to the number of DEGs as the transcriptional efficacy of each ligand (Supplemental Table 1).^50^

**Figure 2.**
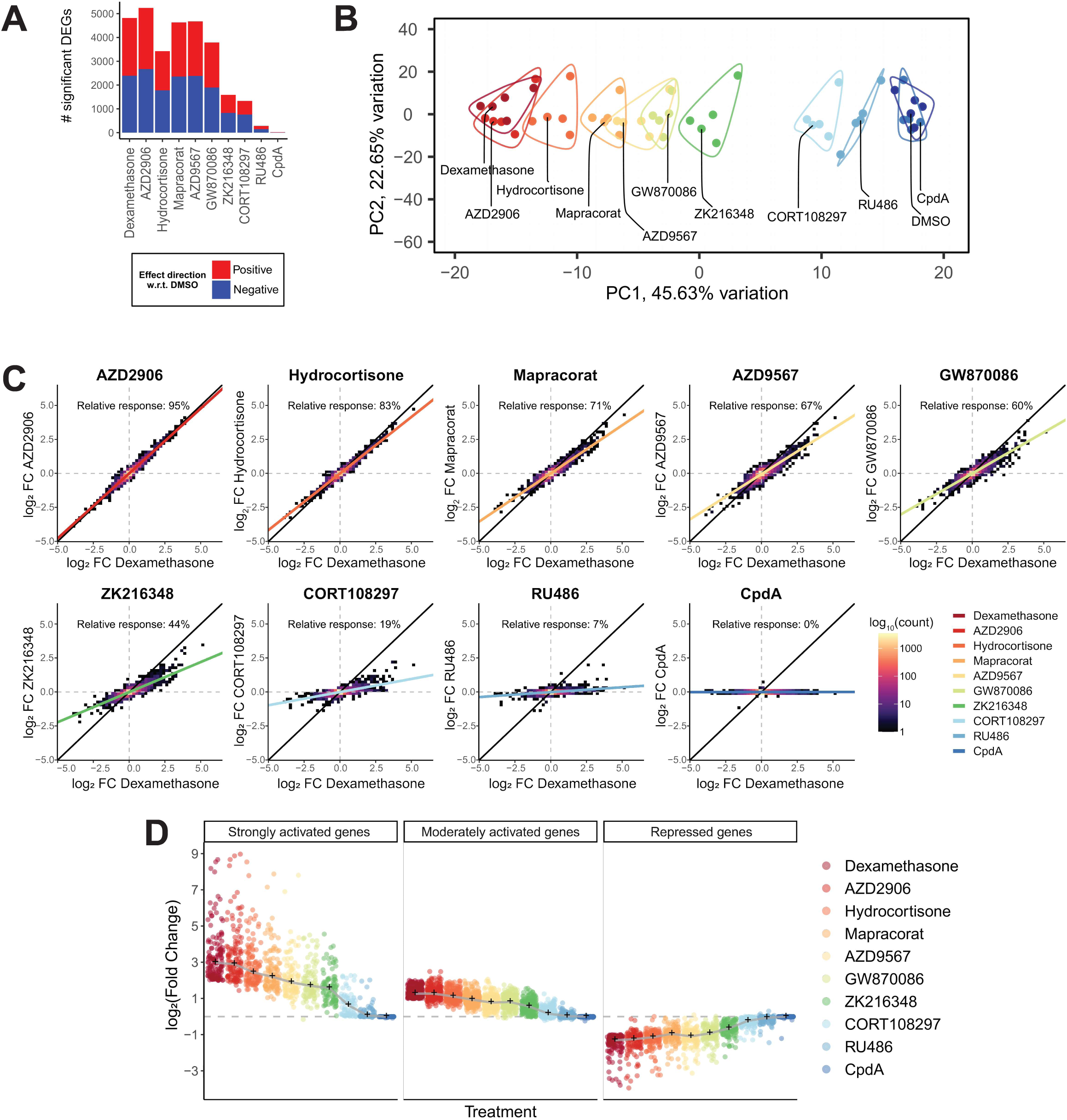
Gene expression responses to each ligand condition vary linearly with respect to dexamethasone-responsive effects. A. Number of significant differentially expressed genes detected for each ligand. B. Principal component analysis of the observed transcriptional responses using the union of all ligand-responsive differentially expressed genes. C. Quantification of the linear relationships between the gene expression effects estimated for each ligand and dexamethasone. The relative response percentages refer to the slope of the line fit by linear regression. D. The magnitude and sign of effect size estimates for strongly and moderately activated genes compared with repressed genes in all conditions. Black markers indicate median effect sizes for each ligand condition.

As expected, GR ligands that induced a greater number of DEGs also induced the larger effects on transcription. Specifically, the transcriptional efficacy correlated with the maximal effect size with a coefficient of R = 0.93 (Fig. S2A). The mean and median effect sizes were more poorly correlated with the number of DEGs (R = 0.57 and R = -0.03, respectively), likely because the mean and median effect sizes are heavily influenced by the number of genes with small effect sizes that meet our criteria for statistical significance. These results indicate that the GR ligands that impact a small fraction of the DEGs shared with dexamethasone had weaker effects on transcription overall.

To identify major modes of the observed transcriptional response, we used principal component analysis applied to all DEGs (Fig. 2B). Principal component 1 (PC1) accounted for 46% of the total variation in gene expression and explained nearly all the GR ligand treatment conditions (R^2^ = 0.96; Fig. S2B). We therefore interpret PC1 as the most ligand-dependent aspect of the glucocorticoid gene expression response. The DEGs with the largest weights from PC1 included well-known GC-responsive genes such as *TFCP2L1*, *CIDEC*, and *FKBP5* (Fig. S2C).^40,51^ The genes in PC1 were also enriched in pathways related to the endocrine system, lipid metabolic processes, and immunity (Fig. S2D). These results were consistent with canonical GC-induced effects on metabolism and the stress response. PC2 accounted for an additional 23% of the total variation in gene expression. PC2 was not explained by variation in GR ligand responses but was explained to some degree by the replicate structure in this study (R^2^ = 0.15).

Based on the above analyses, we hypothesized that the tested GR ligands all activate the same gene expression response, albeit to different degrees. To test that hypothesis, we examined whether a consistent linear relationship existed between the responses to each GR ligand and dexamethasone. We fit linear regression models to the effect sizes estimated for each ligand across all dexamethasone-responsive genes (Fig. 2C). In support of our hypothesis, the average R^2^ across all linear regression models was 0.65 ± 0.35. Moreover, transcriptional effects induced by each GR ligand largely shared the same direction (i.e., activated or repressed), and the maximal effect sizes for each ligand were consistent with the model coefficient. Like our previous analysis, the average R^2^ was significantly and consistently larger in the group of six GR agonists (AZD2906, hydrocortisone, mapracorat, AZD9567, GW870086, and ZK216348; 0.86 ± 0.08) compared to the remaining three ligands (CORT108297, RU386, and CpdA; 0.23 ± 0.24; p = 0.038, Welch’s t-test). We further assessed the fit of each model by estimating the residuals for all conditions. The average residual of 0.093 ± 0.11 was significantly smaller than the average residual for a randomized negative control model (average = 0.21 ± 0.33; p < 2.2×10^-16^, Welch’s t-test). Based on the performance of the linear models, we concluded that the strong agonist response as exemplified by dexamethasone is the core glucocorticoid gene expression response, and the other GR ligands we tested activate the same core response but with varying overall efficacy.

Based on that analysis, we defined the “relative gene expression response” as the slope of the line fit for the gene expression responses between each ligand and dexamethasone. A larger relative gene expression response, represented as percentage close to 100%, indicates that the GR ligand more effectively recapitulates the effects of dexamethasone on gene expression. We observed the largest relative gene expression response of 95% for AZD2906 and the smallest of 0% for CpdA. In a therapeutic context, ligands like RU486 and CpdA likely act as antagonists in competition with cortisol, an endogenous ligand. In the absence of an endogenous agonist, however, these metrics suggest that RU486 (7%) and CpdA induce few or minimal transcriptional effects, whereas SGRMs with either full or partial agonist activities like AZD2906 and mapracorat (71%) induce many more and stronger transcriptional effects like dexamethasone.

We confirm these conclusions by first stratifying the differentially expressed genes by the strength of their response to dexamethasone (i.e., strongly activated, moderately activated, or repressed) and measuring the difference in median ligand-specific effect size (Fig. 2D). We defined strongly activated genes as dexamethasone-responsive genes with at least a four-fold increase in gene expression, moderately activated genes as those with between a two- and four-fold increase in expression, and repressed genes as those with a more than two-fold decrease in expression (Fig. S2E). For example, in response to dexamethasone, strongly activated genes had a median log_2_-fold change of 2.98, whereas the median log_2_-fold change in response to the SGRM agonist ZK216348 was half that magnitude at 1.58. In response to the antagonist CORT108297, the median log_2_-fold change was one-fifth that magnitude at 0.63. This trend was consistent for the moderately activated and repressed genes as well, indicating that each GR ligand effectively modulates a range of dexamethasone-responsive effects, rather than only targeting the most ligand-responsive genes.

### Modulation of regulatory element activities by GR ligands compared to dexamethasone

Based on our observation of a core transcriptional response across all ligands, we hypothesized that there would also be a core response in terms of regulatory element activity. To test that hypothesis, we estimated genome-wide changes in regulatory element activity in response to each ligand using the STARR-seq assays described above (Fig. S3A). We observed between 31,294 and 1,397 gene regulatory element responses across those treatments, and we refer to that number as the regulatory efficacy of each ligand (Fig. 3A, Supplemental Table 2). Overall, the regulatory efficacy was well correlated with the maximal effect size for each ligand (R = 0.79; Fig. S3B). The mean and median regulatory element responses were roughly 3-fold and 4-fold larger, respectively, than the respective transcriptional responses, suggesting a greater dynamic range in regulatory element activity than gene expression responses.

**Figure 3.**
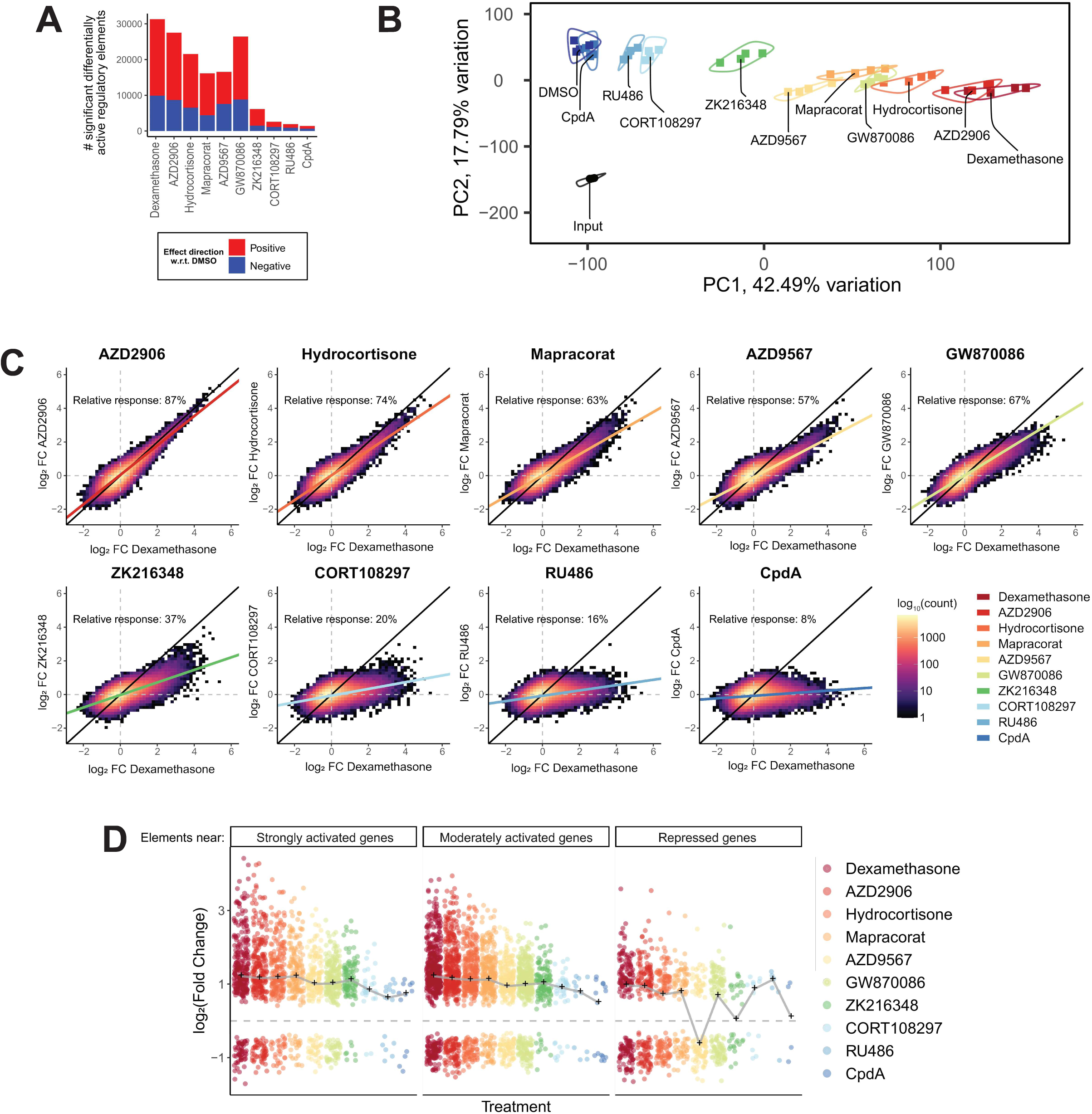
Regulatory element activity responses to each ligand condition are similarly correlated with dexamethasone-responsive effects. A. Number of regulatory elements with significant differential activity detected for each ligand. B. Principal component analysis of the observed regulatory element responses using the union of all elements with ligand-responsive activity. C. Quantification of the linear relationships between the regulatory element effects estimated for each ligand and dexamethasone. The relative response percentages refer to the slope of the line fit by linear regression. D. The magnitude and sign of effect size estimates for regulatory elements near strongly and moderately activated genes compared with element near repressed genes in all conditions. Regulatory elements were annotated to the nearest neighboring gene based on genomic distance. Black markers indicate median effect sizes for each ligand condition.

Variation across ligands in the number of identified regulatory element responses and in the magnitudes of those responses was largely consistent with the trends in gene expression responses reported above. For example, dexamethasone, AZD2906, Hydrocortisone, Mapracorat, AZD9567, and GW870086 had the most numerous and strongest effects on regulatory element activity (mean = 23,242 ± 6,190 ligand-responsive regulatory elements). Most of the observed agonist responses involved regulatory element activation (Supplemental Table 2). We also found that the antagonists CORT108297, RU486, and CpdA induced differential activity at substantially fewer regulatory elements (mean = 3,012 ± 2,144 ligand-responsive regulatory elements). The agonist ZK216348 was an outlier in that the effect sizes and number of elements impacted was on par with the antagonists. Nevertheless, the effects of ZK216348 on regulatory element activity were more highly correlated with the canonical agonist response (R^2^ = 0.64 vs. dexamethasone) than with the antagonist response (average R^2^ = 0.40).

To better define the major modes of the effects of glucocorticoids on regulatory element activity, we again used a principal component analysis applied to all ligand-responsive regulatory elements (Fig. 3B). Principal component 1 (PC1) accounted for 42% of the total variation in regulatory element activity, and GR ligand treatment conditions dominantly explained that response (R^2^ = 0.94; Fig. S3C). Consistent with the results from RNA-seq, we interpret PC1 as a measurement of the most ligand-dependent regulatory element activities. PC2 accounted for an additional 18% of the total variation in regulatory element activity. That component was largely explained by the differences between input and output libraries (R^2^ = 0.81). We therefore interpret PC1 as the core gene regulatory element response across GR-targeting ligands.

We next evaluated if the same linear relationships we identified in the transcriptional responses to the tested GR ligands would also exist in the regulatory element activity responses. Specifically, we hypothesized that the GR ligands largely induce differential activity at the same regulatory elements as dexamethasone, rather than discrete subsets of those elements or different elements entirely. We fit linear regression models to the effect sizes estimated for each ligand across all dexamethasone-responsive regulatory elements (Fig. 3C). The average R^2^ was 0.66 ± 0.32. Again, we found that the regulatory element activity responses to the group of six GR agonists (AZD2906, hydrocortisone, mapracorat, AZD9567, GW870086, and ZK216348) were better correlated (R^2^ = 0.86 ± 0.09) with the linear models than the remaining three antagonists (CORT108297, RU386, and CpdA; R^2^ = 0.26 ± 0.15; p = 0.01, Welch’s t-test). Generally, the STARR-seq data had a wider spread with respect to the linear regression models compared to the RNA-seq data (p = 0.004, Wilcoxon test; Fig. S3D). These correlations were consistent with the previous analysis comparing the ligand-specific gene expression responses with the response to dexamethasone. We also assessed the fit of the regulatory activity response models by measuring the residuals for all conditions. The average residual of 0.22 ± 0.19 was roughly twice as large as the average residual for the RNA-seq linear models but significantly smaller than the average residual for a randomized negative control model (average = 0.47 ± 0.44; p < 2.2×10^-16^, Welch’s t-test). We concluded that the ligands we tested produced regulatory element activity responses that varied linearly with the response to dexamethasone. This behavior was consistent across all STARR-seq regions, not just a subset of elements or elements that were not responsive to dexamethasone as we would expect from a GR ligand that impairs dimerization and/or DNA binding. Therefore, we hypothesized that ligand-specific regulatory element activity responses are tuned according to the ability of the ligand to activate the TF activity of GR - that is, the strength of the agonist activity of the ligand - rather than differential occupancy at GR binding sites.

Based on the above observations, we defined the “relative regulatory element response” following a similar approach as we used to define a relative gene expression response above. Specifically, a larger relative regulatory element activity response, represented as percentage close to 100%, indicates that the response to that GR ligand was more similar to the response to dexamethasone. The same ligands that had large relative gene expression responses also had large relative regulatory element responses, and the same was true for the smaller relative responses (Supplemental Table 3). We observed the largest relative regulatory element activity response of 87% for AZD2906 and the smallest of 8% for CpdA; the remaining ligands had intermediate relative responses. Interestingly, we did not observe a strong correlation between the magnitudes of differential regulatory element activities and differential gene expression, suggesting that the relationship between the two is not trivially modeled (Fig. 3D).

### Explanation of variance associated with SGRM-responsive effects

One approach for developing more effective GC therapies has focused on increasing the affinity of the ligand for GR.^52–54^ Here, though, we find that gene expression and regulatory element activity responses to each ligand were not associated with the affinity for GR. First, we compared the half-maximal inhibitory concentrations (IC_50_) for each ligand to the number of ligand-responsive effects (i.e., efficacy). We also compared the IC_50_ to the relative responses to each ligand, which estimate the degree to which the ligand replicates the effects of dexamethasone. Overall, there were strong correlations between the respective efficacies and relative responses of each ligand for both the gene expression (R^2^ = 0.87) and regulatory element activity (R^2^ = 0.93), suggesting that these measures explain similar properties of the ligands (Fig. 4A). In contrast, there was no significant correlation between those values and the IC_50_ values (Fig. 4B). Of note, a previous study of GR activity in response to RU486 found that the level of partial agonist activity was related to GR concentration and independent of the dose-response curve.^55^ Taken together, we interpret these relationships as a demonstration that, although compounds with higher affinity for GR can be used at lower concentrations, the magnitude of the maximal effects on gene expression and regulatory element activity elicited by each ligand varies independently of that affinity.

**Figure 4.**
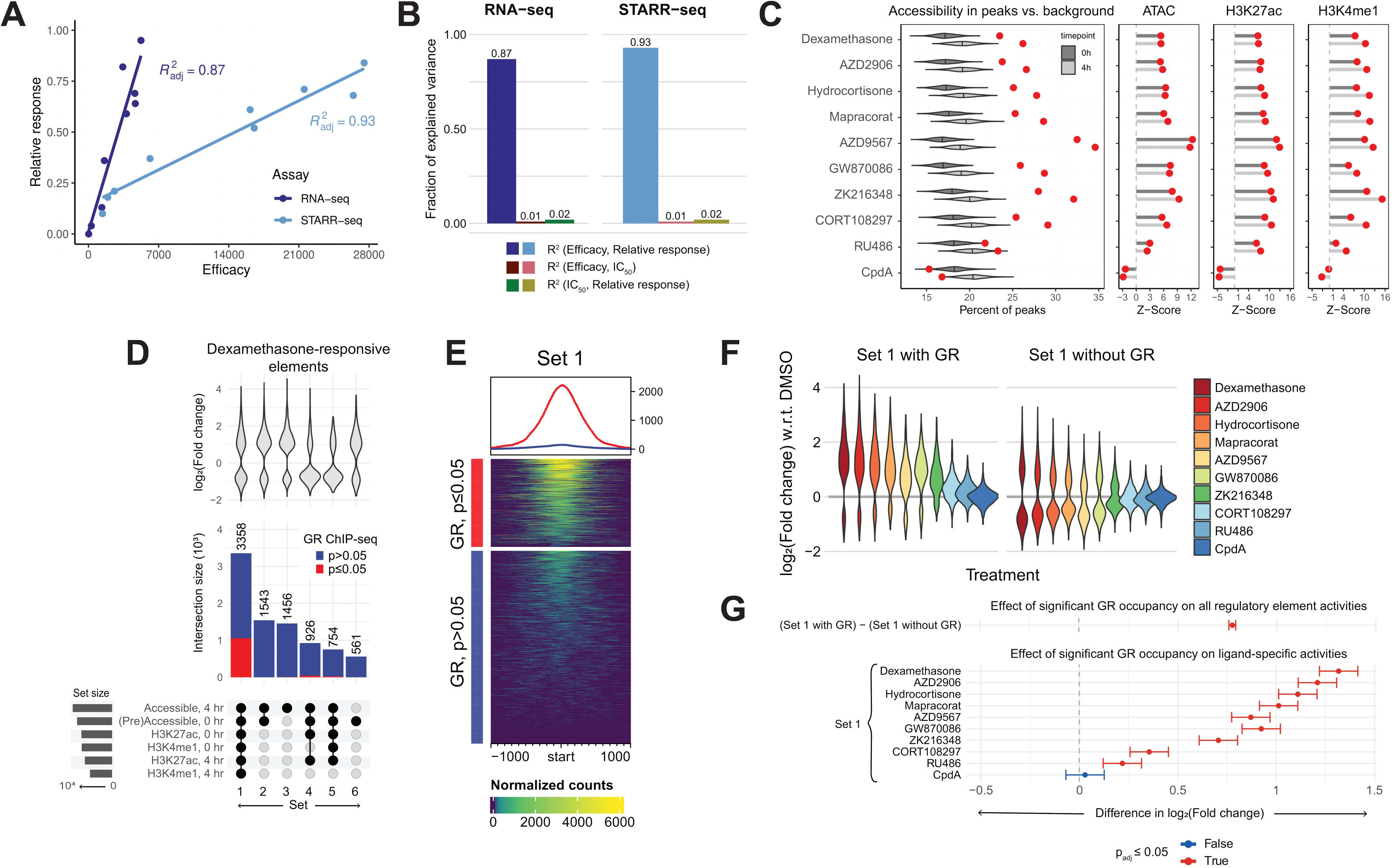
Variation in genomic responses is dominantly explained by the number of ligand-responsive effects, not by affinity, and is associated with chromatin context. A. Correlation from linear regression model fits to the relative responses and efficacies from each assay. B. Correlation from linear regression model fits to the IC50 values for each ligand. C. Percentage of regulatory elements overlapping regions of accessible chromatin and H3K27ac and H3K4me1 histone modifications. The distribution of percentage overlap between background regions and accessible chromatin is shown as a violin plot for each timepoint. The percentage of overlap between ligand-responsive elements and accessible chromatin is indicated with a red circle. Z-scores were calculated based on these measurements. D. The intersection of chromatin markers, GR occupancy, and ligand-specific regulatory element activity. Intersection size for regulatory elements that overlap with regions of the genome marked by accessibility, H3K27ac, H3K4me1, or GR occupancy are shown along with the distribution of dexamethasone-responsive effect sizes for those elements. E. GR occupancy signal measured by ChIP-seq for all regulatory element loci from Set 1, indicated in A. Significant GR occupancy means that a ChIP-seq peak was called with p < 0.05. F. Difference in activities of regulatory elements in Set 1 that do and do not overlap detected GR occupancy sites. G. Confidence internals estimated for differential activity associated with detected GR occupancy in Set 1 regulatory elements using the Tukey HSD test. Confidence internals that span the origin indicate no significant difference in activity associated with GR occupancy.

### Integration of regulatory elements with markers of regulatory chromatin

Where GR binds the genome is influenced by several contextual features, such as the accessibility of the genomic region and the presence of a glucocorticoid receptor DNA binding sequence.^40,49^ Cell- and tissue-specific GR activity additionally arises from the availability of TFs and coregulators that GR interacts with to regulate gene expression. Importantly, upon binding, the ligands that we tested are likely to have conferred specific changes to the conformation of GR, theoretically affecting the dimerization and DNA binding potential of the receptor. We hypothesized that upon activation by these ligands GR may interact (or not interact) with different parts of the genome compared to activation of GR by dexamethasone. To test this hypothesis, we measured whether the regulatory elements with ligand-specific activities that we identified were in genomic regions marked by certain chromatin contexts.

To evaluate relationships between GR binding, chromatin state, and regulatory element activity, we measured the overlap between regulatory elements identified with STARR-seq, regions of chromatin accessibility identified using ATAC-seq, GR binding sites identified with ChIP-seq, and covalent histone modifications associated with active enhancers also identified using ChIP-seq. If the ligands we tested do alter the affinity of GR for binding DNA, we would expect to see differential enrichment of these markers as a reflection of the difference in GR conformational state.^1,27^ For this analysis, we compared our results to that from a previous multi-omic time course analysis of the response to 100 nM dexamethasone.^40^ The genomic response to glucocorticoids was consistent between the two studies. For example, GR has previously been shown to predominantly bind genomic regions where chromatin is accessible prior to glucocorticoid treatment (i.e., pre-accessible).^49^ Consistent with that result, 70% of the GR occupancy sites (N = 3,826) we detected after a 4 hr., 100 nM dexamethasone treatment were in pre-accessible chromatin (Fig. S4A).

We first focused on whether ligand-responsive regulatory elements were similarly enriched in genomic regions with pre-accessible chromatin. In contrast to the extent of GR binding in pre-accessible chromatin, we found that about one-third of the regulatory elements we identified with STARR-seq overlapped with pre-accessible chromatin (maximum = 35% for AZD9567 and minimum = 23% for CpdA). We saw a roughly 3% increase in that overlap with the 31,294 dexamethasone-responsive regulatory elements 4 hr. after dexamethasone treatment. AZD9567 had the largest percentage of ligand-responsive regulatory elements overlapping accessible chromatin at 35%, whereas the remaining agonists ranged between 25-29%. Notably, RU486-responsive regulatory elements had the smallest, though still significant, enrichment over the background regions, and CpdA-responsive regulatory elements had less overlap with accessible chromatin than background regions. Several factors may contribute to that partial overlap. Likely due to its episomal nature, STARR-seq can identify latent regulatory elements – i.e., elements that are in closed chromatin in A549 cells and have potential regulatory activity in a different cell type.^56,57^ It is also possible that many of the GR ligand-responsive STARR-seq activities may be downstream of primary GR activation. Consistent with this reasoning, only 5% of the dexamethasone-responsive regulatory elements in our STARR-seq analysis overlapped significant GR binding sites from ChIP-seq (Fig. S4B).

We next focused on whether the tested ligands selectively activate subsets of regulatory elements marked by specific covalent histone modifications. Indeed, the regulatory elements activated by each ligand have similar histone modification states in their original genomic context (Fig. 4C, Fig. S4C). The most frequent combination of chromatin markers that intersected with dexamethasone-responsive regulatory elements (N = 3,358) had chromatin accessibility as well as H3K27ac and H3K4me1 modifications both before and after dexamethasone treatment (Fig. 4D). GR occupancy was detected at 31.3% of those regions (N = 1,053), which was the greatest enrichment of detected GR occupancy at any combination of chromatin markers (Fig. 4E). We also identified dexamethasone-responsive regulatory elements that solely overlapped with regions of chromatin with accessibility before and/or after dexamethasone treatment and no other contextual markers (Fig. S4D).

Finally, we investigated if there were differences in GR binding that correspond to ligand-specific regulatory element responses (Fig. 4F). Generally, the activities at regulatory elements that overlap a site occupied by GR were larger in magnitude and more positive (1.7-fold) than the activities at regulatory elements that did not overlap a site occupied by GR (Fig. 4G, Fig. S4E). The difference in these activities was largest for the dexamethasone-responsive enhancers with a 2.5-fold increase in activity for elements that overlap a GR occupancy site over those that do not. All other ligands, except for CpdA, had significantly increased activity at elements that overlapped a GR occupancy site, albeit with differences in the magnitude of activity (Fig. 4G, Fig. S4F). However, although we found that regulatory elements do indeed have different levels of activity depending on if they overlap with a region of GR occupancy, the increased activity at these elements suggests that the affinity of GR for DNA binding is not fully dissociated by the ligands we tested.

### Simulation of regulatory element activities under a selective regime

Overall, we demonstrated that the ligands we tested generally activate the same gene expression and regulatory element responses, with the maximal effects varying linearly with respect to dexamethasone. SGRMs, however, were designed to elicit significant responses at a subset of dexamethasone-responsive genes. Theoretically, the genes that are regulated by GR and have anti-inflammatory or immunomodulatory functions would be activated by all GR ligands to maintain the mechanism of action of these therapies. Genes that are not activated by SGRMs would include those normally regulated by GR but with functions unrelated to the control of inflammatory processes. Thus, the SGRMs would not activate genes responsible for the manifestation of adverse - often metabolic-related - side effects. We extended this theory to apply to the activation of regulatory elements with significant activity in response to GR ligand treatments. Specifically, we hypothesized that SGRMs would similarly elicit significant responses at either a subset of dexamethasone-responsive regulatory elements or different elements altogether to achieve dissociation of unwanted side effects.

We developed a simulation to model those scenarios. Specifically, we simulated SGRM responses by selecting a fraction of regulatory element activities to replace with random non-significant activities from elsewhere in the genome (see Methods for details). We then used two-component Gaussian mixture models to investigate if we could identify such selective regimes in our STARR-seq data (Fig. 5A, Fig. S5). Specifically, we expect one component of the model to be a cluster of selectively targeted regulatory elements for which the simulated effects correlate strongly with the observed dexamethasone response, and we expect the second component to consist of non-targeted elements for which the simulated effects have near-zero correlation with the observed dexamethasone response.

**Figure 5.**
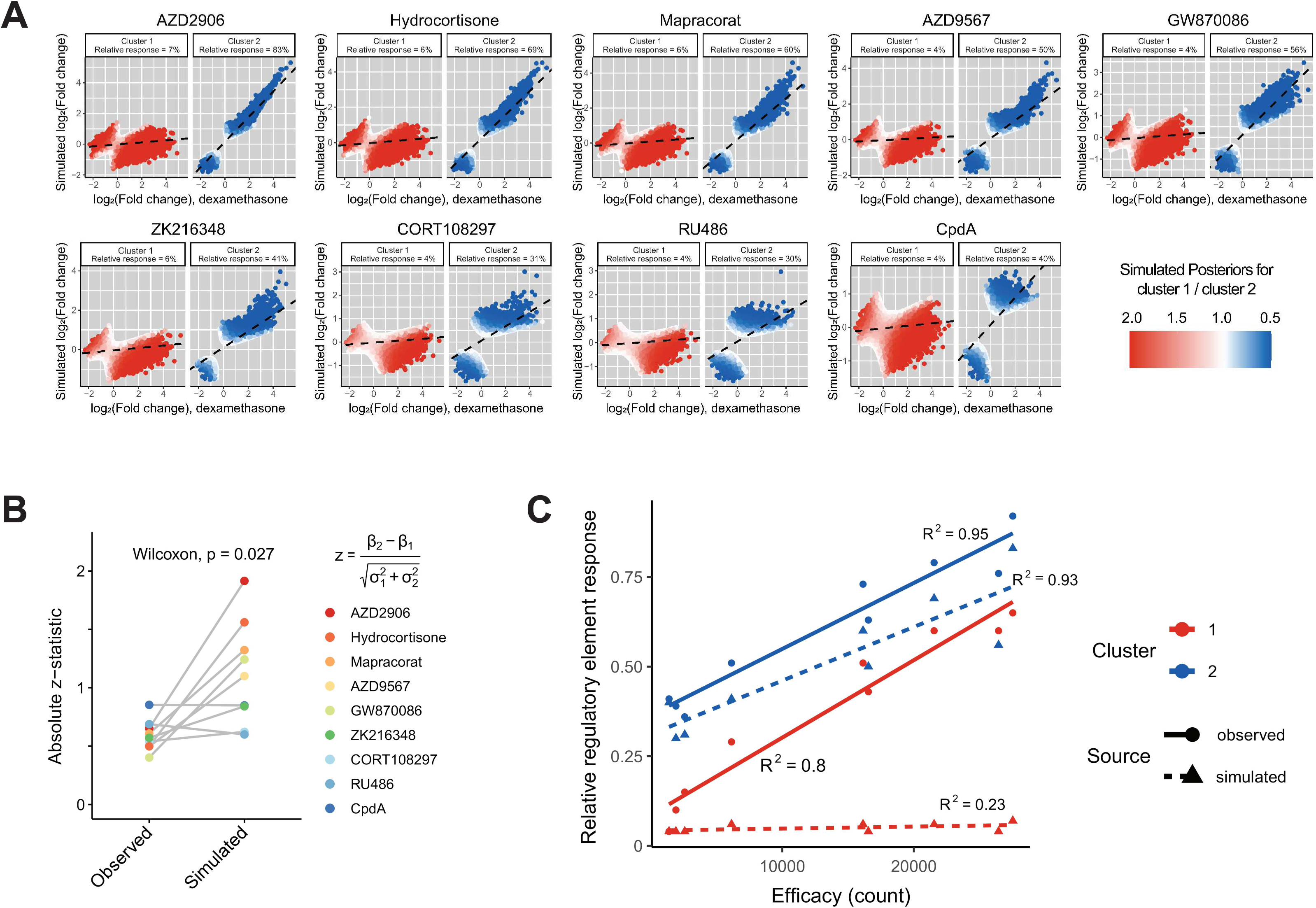
Simulation of regulatory element activation by a hypothetical, highly-selective GR ligand compared to observed responses. A. Simulated regulatory element responses using a two-component Gaussian mixture model for clustering. Elements are assigned to each cluster according to the maximum posterior probability, and relative responses for each cluster are estimated using linear regression. B. Comparison of linear regression models fit to clusters 1 and 2 for each pair of observed and simulated datasets. The model coefficients and standard errors were used to calculate z-statistics which were then used to perform a Wilcoxon test for significance. C. Correlations between the relative responses and regulatory element efficacies for each cluster in paired observed and simulated datasets.

Several of the observed datasets (e.g., AZD2906, hydrocortisone, GW870086) had similar relative response estimates close to 100% in both components, and we interpreted those results as evidence that these ligands do not selectively target a subset of dexamethasone-responsive regulatory elements for activation (Fig. S5). For example, in contrast to our expectations, the AZD2906-responsive regulatory element activities in cluster 1 had a relative response of 65%, and the activities in cluster 2 had a relative response of 92%. Notably, the expected separation between the clusters was present only in the simulated datasets. For example, the simulated AZD2906-responsive regulatory element activities in cluster 1 had a relative response of 7%, whereas cluster 2 had a relative response of 83%. That observation demonstrates we could detect such a selective response were it present (Fig. 5A).

Overall, the results of our simulation experiment suggest that the regulatory element responses we observed in each GR ligand condition were different from the response to a hypothetical, highly selective SGRM. Specifically, we did not find that a subset of dexamethasone-responsive regulatory elements was not targeted for activation in most ligand treatment conditions, except for the antagonists that we tested. The support for this conclusion comes from the fact that relative regulatory responses for elements in cluster 1 of the observed mixture models were not close to the expected 0%. We compared the performance of linear regression models fit to each cluster in the simulated and observed datasets by calculating a z-score based on the model coefficients (Fig. 5B). Indeed, the differences between the relative responses in clusters 1 and 2 of the observed mixture models were significantly smaller than the same differences in the simulated mixture models (p = 0.027, Wilcoxon test). Therefore, we reject the null hypothesis that the observed regulatory element activities and those that we simulated for a hypothetical, highly selective SGRM were the same.

Furthermore, we show that the relative responses for clusters 1 and 2 of the observed mixture model are highly correlated with the regulatory efficacy at R^2^ = 0.80 and R^2^ = 0.95, respectively (Fig. 5C). However, the relative responses for cluster 1 of the simulated mixture model, corresponding to the subset of dexamethasone-responsive regulatory elements not targeted for activation by our hypothetical SGRM, are poorly correlated with regulatory efficacy (R^2^ = 0.23). In comparison, the relative responses for cluster 2 of the simulated mixture model remained highly correlated with regulatory efficacy (R^2^ = 0.93), indicating that the data in this cluster contains similar information as the observed datasets. Therefore, we conclude that the variation in the response to these non-targeted regulatory elements would, instead, be dominantly explained by a still unknown selective mechanism.

## Discussion

In this study, we have assayed changes in human gene regulatory element activity genome-wide across a wide range of GR agonists, partial agonists, and antagonists. Our results reveal that those compounds all modulate the same core set of regulatory elements and genes, albeit with a wide range of maximal effects. That remarkably consistent response suggests that the interactions between each compound and the GR binding pocket trigger essentially the same downstream mechanisms, albeit to a different maximal amount. Even the antagonist RU486, by itself, triggers a substantial subset of that prototypical response. This result suggests that even antagonists, designed to outcompete agonists for the GR binding pocket, can and do weakly activate the GR in the absence of those agonists.

There are two clear limitations to this study. First, studying regulatory element activity outside of the greater chromatin context restricts the gene regulatory mechanisms assayed. Specifically, we and others have shown that STARR-seq identifies a substantial number of latent regulatory elements – i.e., regulatory elements active in other cell types but not accessible to transcription factors in the chromatin of the A549 cells we used. On the one hand, that enables us to assay a greater number of GR ligand-responsive elements than are present in A549 cells alone. On the other hand, that prevents us from identifying chromatin-dependent gene regulatory mechanisms. We partially mitigate that limitation by using longer (1 kb) DNA fragments than in previous high-throughput reporter assay studies.^39,43^ That enables us to better recapitulate interactions between transcription factors at that length scale, and increases the potential for appropriate chromatinization on the plasmids. However, longer-range interactions can and do play an important role in GC responses.^58^ Although that limitation may conceal some ligand-specific regulatory element responses, we note that the overall gene expression responses do depend on that chromatin, and they were also similarly consistent across treatments. A second limitation is that we assayed each ligand by itself, and the effects are likely different in the context of endogenous cortisol, especially for Compound A and RU486. For that reason, we do not interpret our results in terms of how they impact endogenous cortisol responses, but rather as indication of the GC response mechanisms that each ligand elicits by itself.

Together, these results suggest that eliciting a specific subset of the GC response may be more readily achieved by simultaneously targeting other proteins involved in the response. For example, there may also be opportunities to avoid adverse hyperglycemic and insulin resistance outcomes by simultaneously targeting histone deacetylases, interacting transcription factors, or even tissue-specific glucocorticoid synthesis.^59,60^ Similarly, recent approaches have been proposed to develop small molecules targeting steroid receptor coactivators as a novel approach for cancer therapy, targeted immunomodulation, and ameliorating GC impacts on wound healing.^61–63^ These approaches have the potential to rationally inhibit a single adverse side effect or promote a desired outcome based on known pathways. Indeed, that may only be possible by targeting coregulatory proteins necessary for GR activity rather than GR itself. The potential search space for such simultaneous treatments is quite large, with over 300 GR co-regulators identified – suggesting both an enormous unrealized opportunity and a substantial challenge.^1^ Improved identification of functional response elements that are upstream of genes directly or indirectly regulated by GR may be valuable information for the design of such future studies.

## Resource Availability

## Lead Contact

Requests for further information and resources should be directed to and will be fulfilled by the lead contact, Tim Reddy.

## Materials Availability

This study did not generate new unique reagents.

## Data and Code Availability

The data generated and used for this study are available to be downloaded from the ENCODE portal (https://www.encodeproject.org/) with the following identifiers:

Paired mRNA-seq and STARR-seq datasets

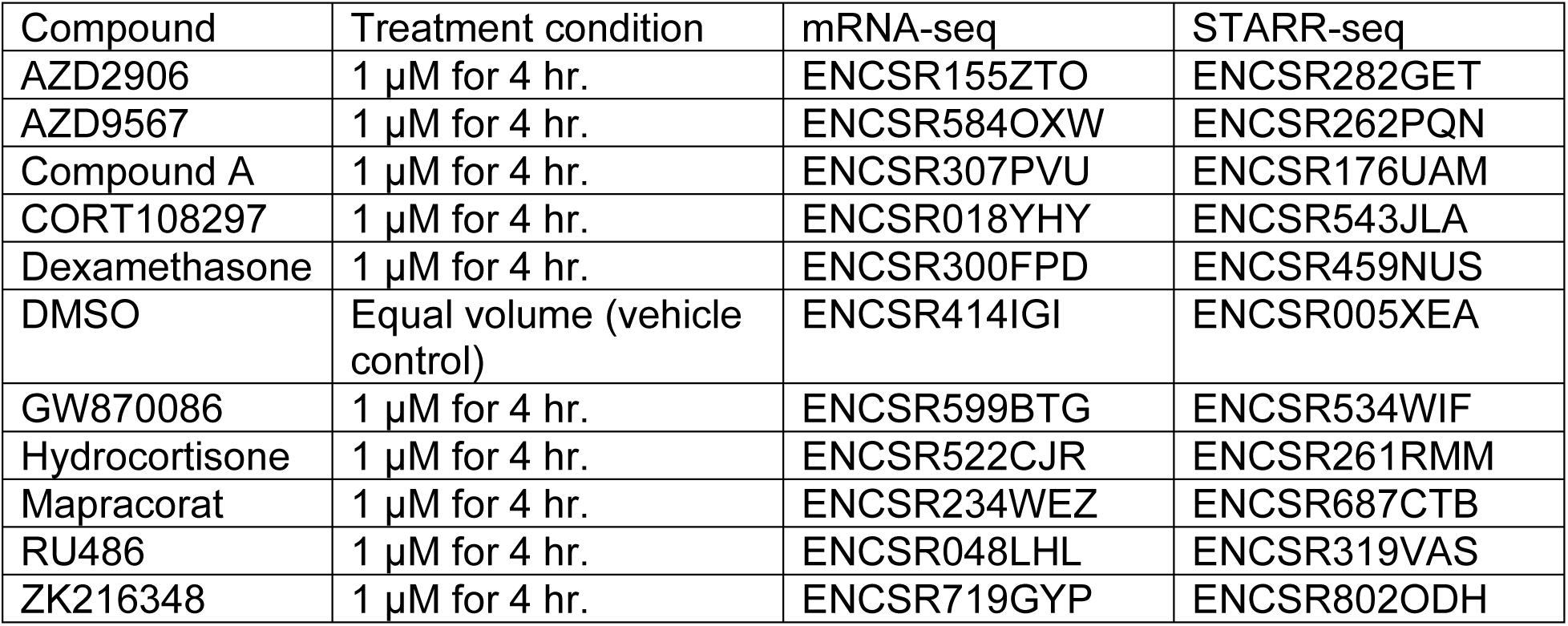

| Compound | Treatment condition | mRNA-seq | STARR-seq |
| --- | --- | --- | --- |
| AZD2906 | 1 μM for 4 hr. | ENCSR155ZTO | ENCSR282GET |
| AZD9567 | 1 μM for 4 hr. | ENCSR584OXW | ENCSR262PQN |
| Compound A | 1 μM for 4 hr. | ENCSR307PVU | ENCSR176UAM |
| CORT108297 | 1 μM for 4 hr. | ENCSR018YHY | ENCSR543JLA |
| Dexamethasone | 1 μM for 4 hr. | ENCSR300FPD | ENCSR459NUS |
| DMSO | Equal volume (vehicle control) | ENCSR414IGI | ENCSR005XEA |
| GW870086 | 1 μM for 4 hr. | ENCSR599BTG | ENCSR534WIF |
| Hydrocortisone | 1 μM for 4 hr. | ENCSR522CJR | ENCSR261RMM |
| Mapracorat | 1 μM for 4 hr. | ENCSR234WEZ | ENCSR687CTB |
| RU486 | 1 μM for 4 hr. | ENCSR048LHL | ENCSR319VAS |
| ZK216348 | 1 μM for 4 hr. | ENCSR719GYP | ENCSR802ODH |

The methods for generating these datasets are also available through the ENCODE portal for each experiment. We have reproduced these methods for RNA extraction, RNA-seq, and whole genome STARR-seq here.

| Description | ENCODE URL |
| --- | --- |
| PolyA plus library construction protocol | https://www.encodeproject.org/documents/e4e1b09a-8fe8-4212-9e76-a2fd6d0c8b49/ |
| Transfection protocol for whole genome STARR-seq libraries | https://www.encodeproject.org/documents/c2499c10-935d-4bc9-b374-5a40f69d4af9/ |
| Whole genome STARR-seq protocol | https://www.encodeproject.org/documents/c8891369-b036-4be1-b881-76ca3911151e/ |
| STARR-seq output library protocol | https://www.encodeproject.org/documents/9113e79f-1a7a-4620-bdb7-3b6c7b2c259f/ |

All original code will be deposited at Zenodo and will be publicly available as of the date of publication.

Any additional information required to reanalyze the data reported in this paper is available from the lead contact upon request.

## Acknowledgements

This work was supported in part by NIH grants HG010741, HG011123, and HG009428 (to T.E.R.). Figure 1A was created in BioRender (https://BioRender.com/snfu4fx).

## Author Contributions

Conceptualization, N.S.G., G.D.J., and T.E.R.; Formal analysis, N.S.G., G.D.J., and A.E.B.; Funding acquisition, T.E.R.; Investigation, G.D.J.; Project Administration, T.E.R.; Resources, G.D.J., and T.E.R.; Software, N.S.G., G.D.J., and A.E.B.; Visualization, N.S.G.; Writing – original draft, N.S.G., and T.E.R.; Writing – review & editing, N.S.G., G.D.J., A.E.B., and T.E.R.

## Declaration of interests

The authors declare no competing interests.

## Supplemental Information

Document S1. Figures S1-5 and Tables S1-3.

## Methods

### RNA extraction

Total RNA was extracted from A549 cells 4 hr. following treatment with one of 10 GR ligands or DMSO. All ligands were dissolved in DMSO, and we therefore used equal volume DMSO as a vehicle control. Total RNA was recovered from cell lysates with the Qiagen RNeasy Midi kit including the on-column DNase I digestion step. Eluates were treated with 1 μL RNase Block (Aligent) and stored at -80°C. The NanoDrop (Thermo Fisher Scientific) was used to assess the quantity and purity of the RNA.

### Plasmid and whole-genome STARR-seq input library construction

Total genomic DNA was extracted from lymphoblastoid cells (GM12878) with the Qiagen DNeasy Blood and Tissue Kit and sonicated with the Covaris S2 system to ∼1 kb. Sonicated DNAs were purified with a 0.7x volume of SPRI beads. In 96 separate reactions, 5 µg of sonicated DNA was end-repaired, A-tailed, and ligated to annealed Illumina TruSeq Universal Adapters using the NEBNext DNA Library Prep Master Mix Kit. Adapted DNAs were purified with a 0.7x SPRI beads and eluted in 65 µl of water. Eluted DNAs were enriched via eight cycles of PCR using the Kapa Hifi HotStart system with GC buffer and STARR-seq adaptor primers. PCR was performed under the following conditions: 98°C for 3 min, followed by eight cycles of 98°C for 20 s, 67°C for 15 s, 72 °C for 1.5 min, with a final extension at 72 °C for 5 min. Amplified inserts were purified with a 0.7x volume of SPRI beads and eluted in 65 µl of water. Following purification, the library was gel size selected using Sybr Safe DNA stain and the GeneJET Gel Extraction Kit (Thermo Fisher Scientific). The STARR-seq ORI plasmid (Addgene #99296) was linearized by digestion with AgeI-HF and SalI-HF (NEB) and gel size selected as above. A total of 155 µg of insert library was cloned into the linearized vector using NEBuilder reagents (NEB). A total of 66 individual assembly reactions were performed using a 2:1 insert to vector ratio and a total of 5 pmol of DNA per reaction. After incubating the assembly reactions at 50°C for 1 hour the DNAs were purified twice with 0.7x volumes of SPRI beads, eluted in water, and stored at -20°C. E.Cloni 10G SUPREME electrocompetent cells (Lucigen) were grown from commercial stocks in super broth following standard protocols. Immediately following sequential washes with 10% glycerol, the competent cells were aliquot into 48 0.2 cm electroporation cuvettes and transformed with 4.5 µg of assembled DNA. Following recovery for 1 h in SOC medium while shaking (225 rpm, 37°C), transformations were pooled four to a flask in 1 L of Luria Broth and then grown for 14 h under carbenicillin selection while shaking (225 rpm, 37°C). Reporter input libraries were purified using the NucleoBond PC 10000 EF Giga kit (Macherey-Nagel). Recovered plasmid libraries were pooled prior to aliquoting and storage at -20°C. The quality and diversity of the pooled plasmid library was assessed by sequencing on an Illumina MiSeq. The absence of bacterial genomic DNA and endotoxins were confirmed via gel electrophoresis and the ToxinSensor Chromogenic LAL Endotoxin Assay Kit (GenScript), respectively.

### Transfection

A549 cells (first obtained at passage 87) were expanded under standard culture conditions using Ham’s F-12K (Kaighn’s) Medium, 10% v/v FBS, 1% v/v penicillin-streptomycin for a total of seven passages. Cells were collected by trypsinization, pelleted, and resuspended in supplemented Lonza SF electroporation buffer. The whole genome STARR-seq library was electroporated into approximately 109 cells using the Lonza 4D-Nucleofector LV unit and the CM-130 program. Electroporated cells were incubated at room temperature for ten minutes before adding warm antibiotic-free media. Fifty million transfected cells were plated into eleven 500 cm2 plates and allowed to recover in the incubator under standard conditions. Two hours post-electroporation, 20 µl of treatment compound was added to a plate for a final concentration of 1 µM. Four hours post-treatment, cells were rinsed with PBS (pH 7.4), and lysed with 10 mL of RLT buffer (QIAGEN) supplemented with 2-mercaptoethanol (Sigma). Lysates were harvested from plates and passed through a 18-gauge needle twenty times and stored at -80°C. The transfection protocol was repeated four times. Treatments from each transfection are considered biological replicates.

### mRNA-seq and whole-genome STARR-seq output library construction

75 μg of total RNA was subjected to poly-A RNA isolation with Dynabead Oligo-dT25 beads (Thermo Fisher Scientific) according to the manufacturer’s recommended protocol. Poly-A RNAs were treated with Turbo DNase (2 U; Thermo Fisher Scientific) and 1 μL RNase Block at 37°C for 30 min before halting the reaction with the DNase Inactivation Reagent. RNA was reverse transcribed for 2 hr at 50°C with a UMI-containing (8mer) reporter-specific primer using the SuperScript III system (800 U; Thermo Fisher Scientific). In parallel, aliquots of each Poly-A RNA sample were used to template no-enzyme negative control reactions. Following enzyme inactivation, reactions were treated with recombinant RNase A (Sigma) at 37°C for 1 hr, purified with SPRI beads (2.0X), and split across seven wells. The cDNAs were then PCR enriched with primers encoding a sample barcode and Illumina flowcell sequences under the following conditions: 95°C for 3 min; followed by 5 cycles of 98°C for 20 s, 63°C for 15 s, 72°C for 90 s. To determine the number of additional PCR cycles needed to enrich a sample, 5 μL aliquots from a representative reaction from each sample were used to template qPCRs. The original reactions were left on ice during the qPCR protocol. The number of additional PCR cycles required was calculated by subtracting 1 from the number of cycles needed to reach 1/3 of maximum optical signal after 40 cycles of enrichment. The original plates were returned to the thermocycler and subjected to further PCR cycles followed by a final extension at 72°C for 5 min. Products of the negative control reverse-transcription reactions were used to template individual PCRs that were subjected to the same reaction conditions as their corresponding samples. Products from the seven PCRs were pooled and purified with SPRI beads (0.5X). Pooled products and corresponding negative controls were assessed on the Agilent TapeStation prior to sequencing on the Illumina NovaSeq platform.

### mRNA-seq data processing and statistical analysis

mRNA-seq libraries were sequenced as 100 bp paired-end reads on the NovaSeq 6000 platform. After being demultiplexed, the fastq files were used as input to the standard ENCODE bulk RNA-seq processing pipeline. Briefly, quality of the reads was assessed with FASTQC. Adapter sequences were trimmed from the reads using Trimmomatic. The reads were mapped to the hg38 human reference genome using 2-pass alignment with STAR. Specifically, the --sjdbOverhang option was used and set to 49. BAM files were generated with samtools, then quantification of read counts was performed at the gene level using featureCounts.

Differentially expressed genes were identified using DESeq2. Using the gene-level count matrix, contrasts were defined as the comparison between each ligand treatment condition (after 4 hr.) and the vehicle control, DMSO, treatment condition. Benjamini-Hochberg multiple hypothesis correction was applied to estimates of statistical significance to achieve a 5% false discovery rate.

### Peak calling, STARR-seq activity estimation, and statistical analysis

STARR-seq input and output libraries were aligned to the human genome (hg38) with Bowtie2 (version 2.2.4), using the following parameters: bowtie2 -X 2500 -- sensitive. Only properly paired alignments with a MAPQ score ≥ 10 that corresponded to fragments greater than 750 bp in length and aligned outside centromeres, gaps, and blacklist regions were retained in downstream analyses. UMI tags were appended to output alignments with fgbio AnnotateBamWithUmis (version 0.4.0). Tagged alignments were deduplicated with picard-tools MarkDuplicates (version 2.4.1). Following deduplication, the four technical replicate libraries comprising each output biological replicate were merged. DeepTools and the UCSC command line bioinformatic utilities were used to convert between the alignment files and signals files. Visualization of the STARR-seq signal as locus plots was done with the rtrackplot package and as heatmaps with the EnrichedHeatmap package.

Peak calling was performed by running CRADLE separately for each replicate. All input signal files were used as background (‘-ctrlbw’) for each individual output signal file replicate (‘-expbw’). The following parameters were used: -p 52, -pl 200. Otherwise, the default CRADLE options were used as previously described. Peaks that were identified in at least two replicates were retained for statistical modeling. Peaks that were identified within 50 bp of another peak (‘min.gapwidth’) were merged together using the R package GenomicRanges. The multiBigwigSummary utility from deepTools was then used to summarize the counts across all treatment conditions for the union set of significant peaks. We then identified STARR-seq regions with differential activity using the R package limma (version 3.54.1). Contrasts were defined as the comparison between the output library reads measured for each ligand treatment condition and the output library reads measured for the DMSO (vehicle control) condition. Benjamini-Hochberg multiple hypothesis correction was applied to estimates of statistical significance to achieve a 5% false discovery rate. Principal component analysis was performed with the PCAtools package. Linear regression models used the lm implementation available in the stats package.

### Comparisons to other genomic datasets

We compared the ligand-specific activities measured by STARR-seq to regions of the genome with other regulatory markers assayed by ChIP-seq and ATAC-seq. Briefly, A549 cells were treated with 100 nM dexamethasone for 4 hr. Both ChIP-seq and ATAC-seq were performed before treatment (0 hr.) and after treatment (4 hr.). The ChIP targets included the histone modifications H3K27ac, H3K4me1, and the GR. These data were released publicly through ENCODE.

## ATAC-seq and ChIP-seq datasets

| Treatment condition | ATAC-seq | H3K27ac ChIP-seq | H3K4me1 ChIP-seq | GR (NR3C1) ChIP-seq |
| --- | --- | --- | --- | --- |
| 100 nM Dexamethasone for 0 hr. | ENCSR220ASC | ENCSR783SNV | ENCSR636PIN | ENCSR516IVY |
| 100 nM Dexamethasone for 4 hr. | ENCSR288YMH | ENCSR543ZVZ | ENCSR949IDI | ENCSR660RYY |

For this analysis, the peaks identified with ChIP-seq, ATAC-seq, and STARR-seq were resized to 2 kb to account for differences in the library fragment sizes. ChIP-seq data were reported as input-subtracted read density (RPKM). Intersections between STARR-seq regions with regulatory activity and other genomic datasets were determined using the findOverlaps function from the IRanges package. For the analysis comparing the overlap of significant ligand-responsive regulatory elements to background regions, 1,000 STARR-seq regions with no significant ligand-responsive activity were sampled randomly for each of 1,000 iterations. These background regions were used to calculate the z-scores for this analysis.

### Simulation of selective regulatory element activities

For the analysis of the simulated regulatory element activities, we assumed that our hypothetical highly selective SGRM may target just 25% of the dexamethasone responsive regulatory elements for activation. We also assumed that the relative response – i.e., the ability of each ligand to replicate the magnitude of the response to dexamethasone – would be consistent with what we observed. Therefore, 25% of the dexamethasone-responsive elements (N = 7,824) were assigned the activities we measured using STARR-seq for each ligand condition. The remaining 75% of dexamethasone-responsive elements were assigned activities representative of no significant response (i.e., “control-like” effects). These control-like effects were drawn from the response we measured in the CpdA condition which had a relative response close to 0%, indicating little difference in regulatory activity compared to the DMSO baseline. Briefly, sampling was performed by weighting the probability of an element getting chosen by the -log10 adjusted p-value estimated in the dexamethasone treatment condition. The ligand-specific effect sizes of the selected STARR-seq regions were replaced with the corresponding effect sizes that we measured for these regions in the CpdA treatment condition.

These simulated datasets were clustered into two distributions using a Gaussian mixture model. Specifically, the flexmix package was used to compare the simulated ligand-specific effect sizes and the observed dexamethasone effect sizes. The number of components for the mixture model was set as 2. Membership of each element in either cluster 1 or cluster 2 was determined by maximizing the posterior estimates from the flexmix model. Linear regression models were fit to the data in each cluster separately. The models fit to the observed data and the simulated data were compared by calculating the z-score from the model coefficients.

## Notes

### Competing Interest Statement

The authors have declared no competing interest.

