## Supplementary material for "The gene regulatory effects of selective glucocorticoid receptor ligands": Document S1

Table S1. Transcriptional efficacies for all tested ligands.

| Compound | Transcriptional efficacy | # Positive / Negative |
|---|---|---|
| Dexamethasone | 4,818 | 2,426 / 2,392 |
| AZD2906 | 5,240 | 2,569 / 2,671 |
| Hydrocortisone | 3,424 | 1,646 / 1,778 |
| Mapracorat | 4,631 | 2,274 / 2,357 |
| AZD9567 | 4,674 | 2,290 / 2,384 |
| GW870086 | 3,787 | 1,891 / 1,896 |
| ZK216348 | 1,588 | 757 / 831 |
| CORT108297 | 1,333 | 571 / 762 |
| RU486 | 283 | 135 / 148 |
| CpdA | 21 | 11 / 10 |

We defined significant ligand-responsive differentially expressed genes as having an adjusted p-value ≤ 0.05 compared to the negative control DMSO condition. Positive and negative effects were reported with respect to DMSO.

Table S2. Regulatory efficacies for all tested ligands.

| Compound | Regulatory efficacy | # Positive / Negative |
|---|---|---|
| Dexamethasone | 31,294 | 21,396 / 9,898 |
| AZD2906 | 27,513 | 18,844 / 8,669 |
| Hydrocortisone | 21,535 | 14,999 / 6,536 |
| Mapracorat | 16,136 | 11,718 / 4,418 |
| AZD9567 | 16,541 | 8,963 / 7,578 |
| GW870086 | 26,431 | 1,7601 / 8,830 |
| ZK216348 | 6,144 | 4,644 / 1,500 |
| CORT108297 | 2,588 | 1,449 / 1,139 |
| RU486 | 1,919 | 1,002 / 917 |
| CpdA | 1,397 | 732 / 665 |

We defined regulatory elements with significant ligand-responsive differential activity as having an adjusted p-value ≤ 0.05 compared to the negative control DMSO condition. Positive and negative effects were reported with respect to DMSO.

Table S3. Relative gene expression and regulatory element activity response estimates for all ligand conditions compared to dexamethasone.

| Compound | Relative gene expression response | Relative regulatory element activity response |
|---|---|---|
| Dexamethasone | 100% | 100% |
| AZD2906 | 96% | 87% |
| Hydrocortisone | 83% | 74% |
| Mapracorat | 69% | 63% |
| AZD9567 | 65% | 57% |
| GW870086 | 60% | 67% |
| ZK216348 | 39% | 37% |
| CORT108297 | 14% | 20% |
| RU486 | 4% | 16% |
| CpdA | 0% | 8% |

Relative responses correspond to the linear regression model coefficients fit to each distribution of effect sizes in Fig. 2C and Fig. 3C.

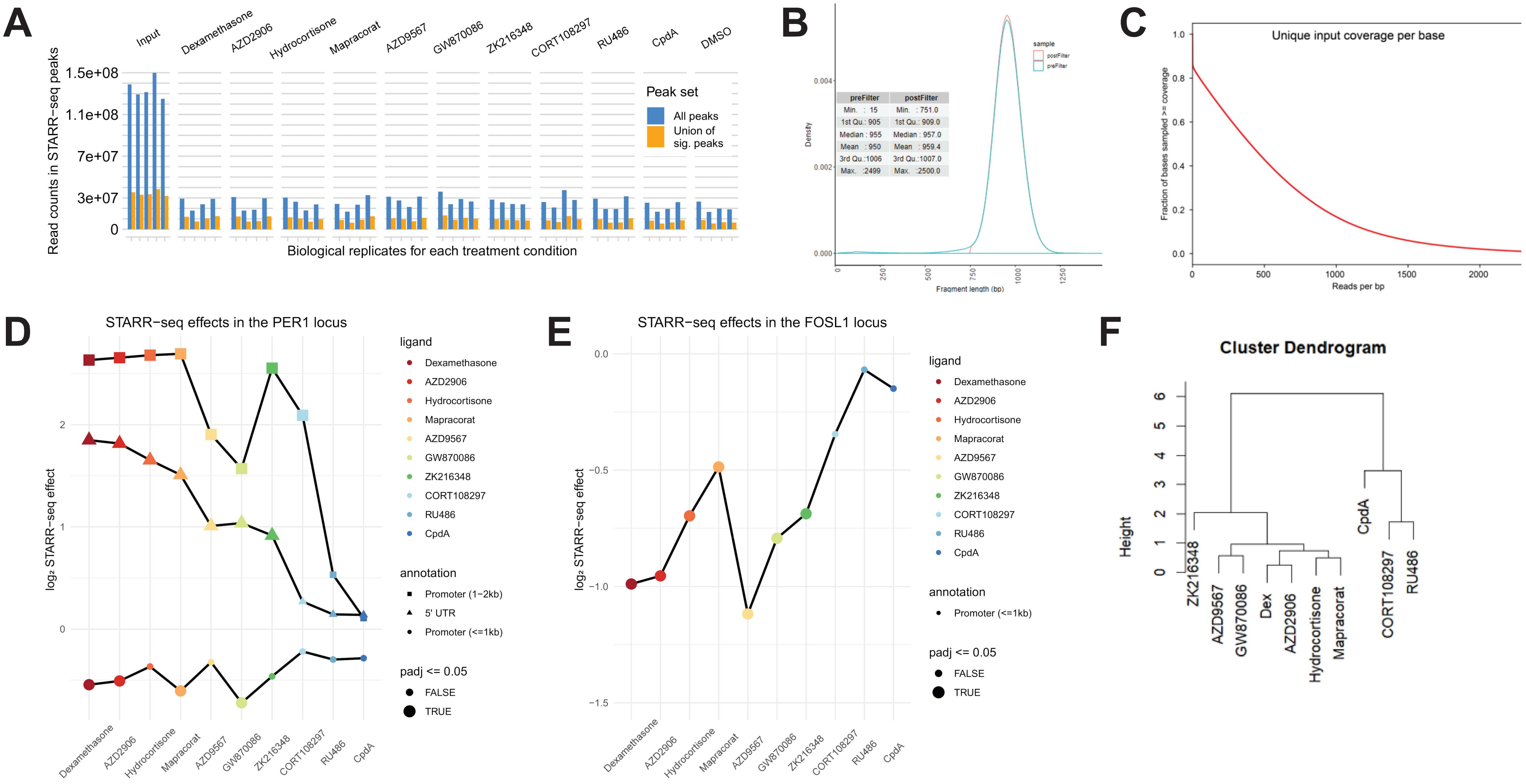
A
Input
Dexamethasone
AZD2906
Hydrocortisone
Mapracorat
AZD9567
GW870086
ZK216348
CORT108297
RU486
CpdA
DMSO
1.5e+08
1.1e+08
7e+07
3e+07
0
Read counts in STARR-seq peaks
Biological replicates for each treatment condition
Peak set
All peaks
Union of sig. peaks
B
sample
postFilter
preFilter
Density
0.004
0.002
0.000
preFilter
Min. : 15
1st Qu.: 905
Median : 955
Mean : 950
3rd Qu.:1006
Max. :2499
postFilter
Min. : 751.0
1st Qu.: 909.0
Median : 957.0
Mean : 959.4
3rd Qu.:1007.0
Max. :2500.0
0
250
500
750
1000
1250
Fragment length (bp)
C
Unique input coverage per base
Fraction of bases sampled >= coverage
1.0
0.8
0.6
0.4
0.2
0.0
0
500
1000
1500
2000
Reads per bp
D
STARR−seq effects in the PER1 locus
log₂ STARR−seq effect
2
1
0
ligand
Dexamethasone
AZD2906
Hydrocortisone
Mapracorat
AZD9567
GW870086
ZK216348
CORT108297
RU486
CpdA
annotation
Promoter (1−2kb)
5' UTR
Promoter (<=1kb)
padj <= 0.05
FALSE
TRUE
E
STARR−seq effects in the FOSL1 locus
log₂ STARR−seq effect
0.0
−0.5
−1.0
−1.5
ligand
Dexamethasone
AZD2906
Hydrocortisone
Mapracorat
AZD9567
GW870086
ZK216348
CORT108297
RU486
CpdA
annotation
Promoter (<=1kb)
padj <= 0.05
FALSE
TRUE
F
Cluster Dendrogram
Height
0
1
2
3
4
5
6
ZK216348
AZD9567
GW870086
Dex
AZD2906
Hydrocortisone
Mapracorat
CpdA
CORT108297
RU486

Figure S1. STARR-seq library characterization, related to Figure 1. A. Number of sequencing reads for each STARR-seq library in all peaks and in peaks with significant activity. B. Distribution of fragments lengths in long-fragment STARR-seq libraries. C. Per-base input coverage across STARR-seq experiments. D. Ligand-specific activities of regulatory elements at the *PER1* locus. E. Ligand-specific activities of regulatory elements at the *FOSL1* locus. F. Hierarchical clustering of the correlations between regulatory element activities in Figure 1.

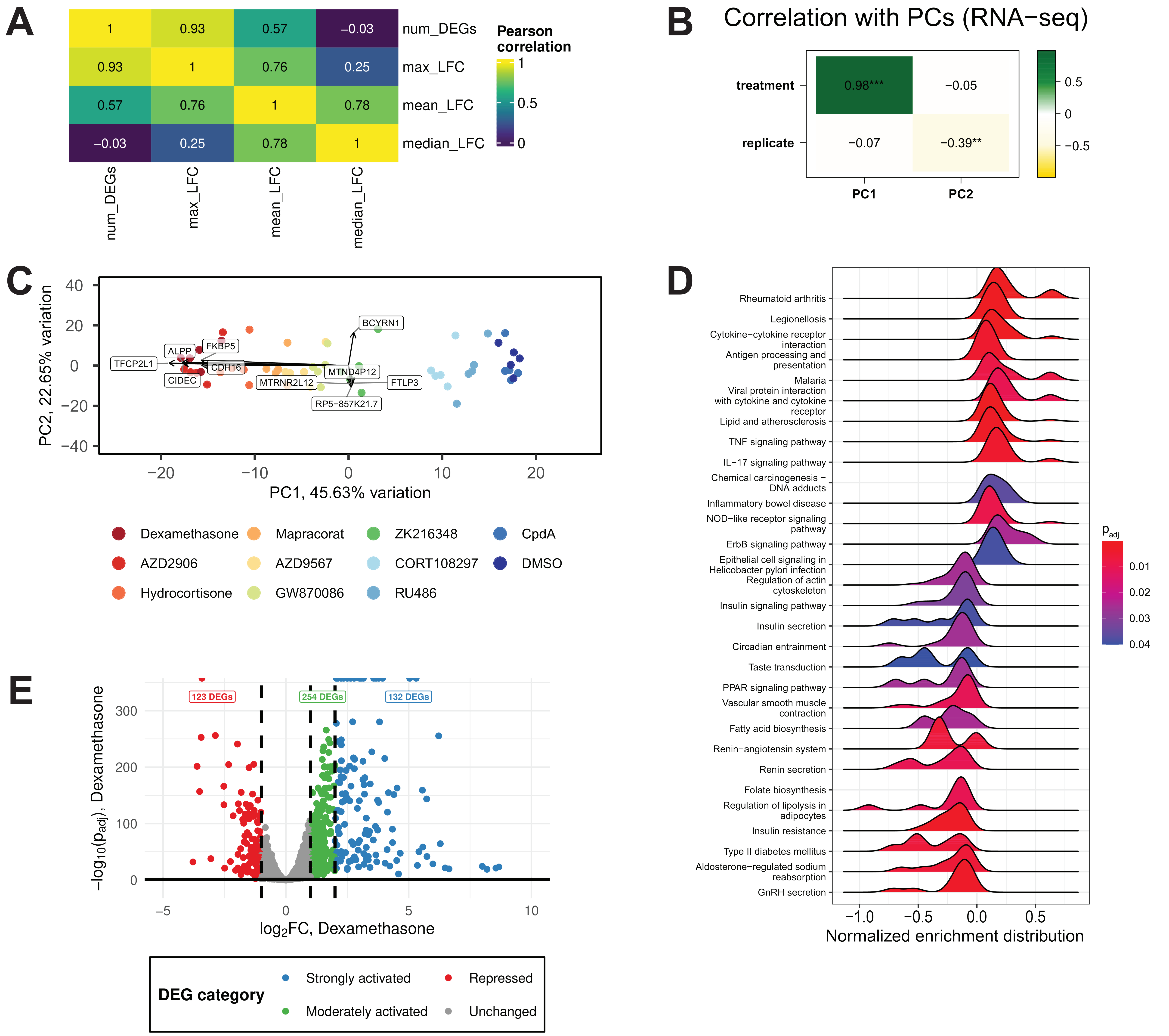
A
num_DEGs
max_LFC
mean_LFC
median_LFC
Pearson correlation
1
0.5
0
0.93
0.57
−0.03
0.76
0.25
0.78
B
Correlation with PCs (RNA−seq)
treatment
replicate
0.98***
−0.05
−0.07
−0.39**
PC1
PC2
0.5
0
−0.5
C
PC2, 22.65% variation
PC1, 45.63% variation
40
20
0
−20
−40
−10
10
BCYRN1
ALPP
FKBP5
TFCP2L1
CDH16
CIDEC
MTND4P12
MTRNR2L12
FTLP3
RP5−857K21.7
Dexamethasone
Mapracorat
ZK216348
CpdA
AZD2906
AZD9567
CORT108297
DMSO
Hydrocortisone
GW870086
RU486
D
Rheumatoid arthritis
Legionellosis
Cytokine−cytokine receptor interaction
Antigen processing and presentation
Malaria
Viral protein interaction with cytokine and cytokine receptor
Lipid and atherosclerosis
TNF signaling pathway
IL−17 signaling pathway
Chemical carcinogenesis − DNA adducts
Inflammatory bowel disease
NOD−like receptor signaling pathway
ErbB signaling pathway
Epithelial cell signaling in Helicobacter pylori infection
Regulation of actin cytoskeleton
Insulin signaling pathway
Insulin secretion
Circadian entrainment
Taste transduction
PPAR signaling pathway
Vascular smooth muscle contraction
Fatty acid biosynthesis
Renin−angiotensin system
Renin secretion
Folate biosynthesis
Regulation of lipolysis in adipocytes
Insulin resistance
Type II diabetes mellitus
Aldosterone−regulated sodium reabsorption
GnRH secretion
p adj
0.01
0.02
0.03
0.04
−1.0
−0.5
0.0
Normalized enrichment distribution
E
123 DEGs
254 DEGs
132 DEGs
300
200
100
−5
5
−log10(padj), Dexamethasone
log2FC, Dexamethasone
DEG category
Strongly activated
Repressed
Moderately activated
Unchanged

Figure S2. Performance of differentially expressed genes for regression and principal component analysis, related to Figure 2. A. Correlation between transcriptional efficacy and log fold-change (LFC) effect sizes for differentially expressed genes. B. Correlations between principal components and experimental variables. C. PCA biplot showing the 10 top loadings in PC1 and PC2. D. Ridge plot showing normalized enrichment scores of top PC loadings in biological pathways. Statistical significance is indicated by color. E. Example categorization of dexamethasone-responsive genes by effect size.

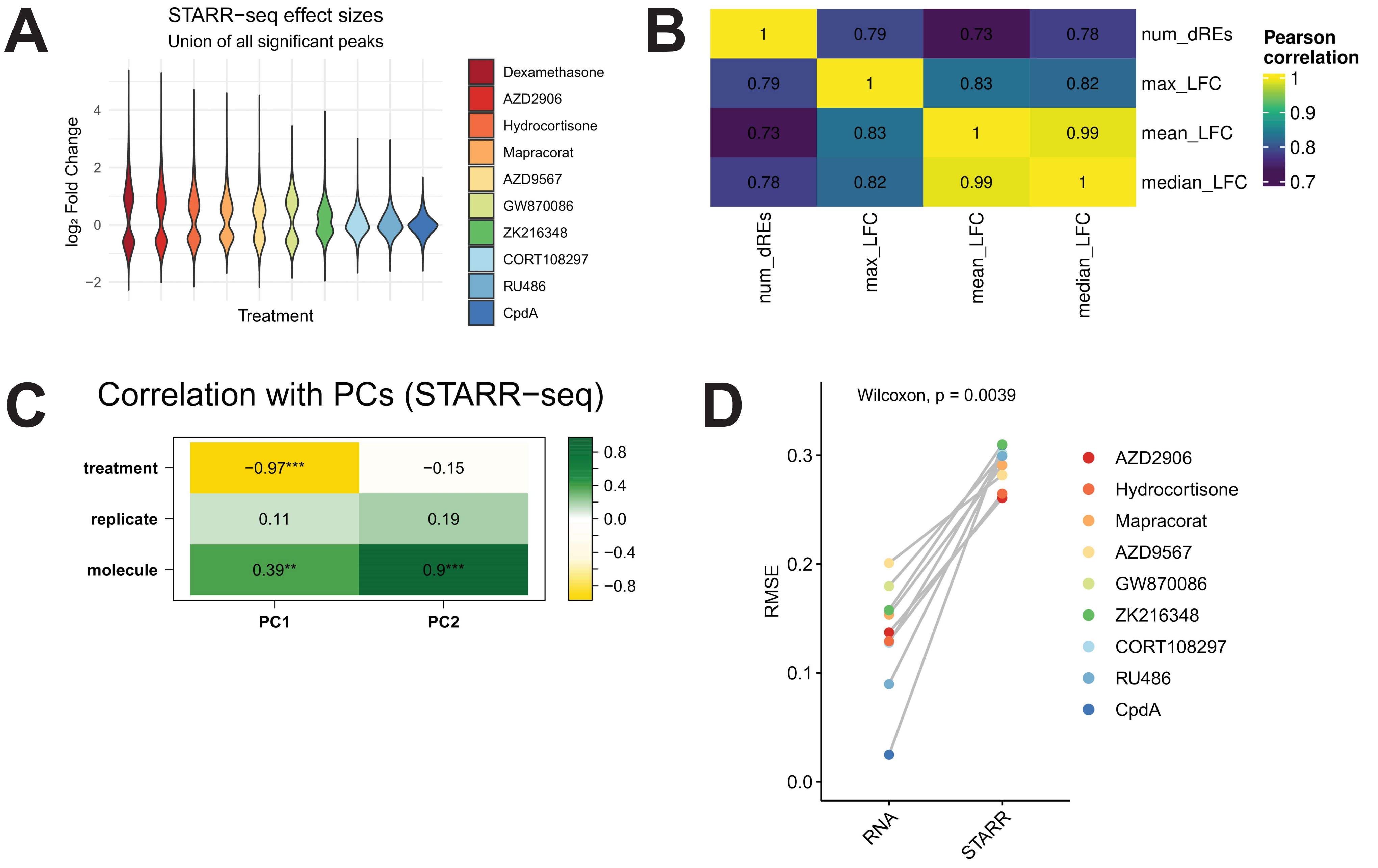

A
STARR−seq effect sizes
Union of all significant peaks
log₂ Fold Change
4
2
0
−2
Treatment
Dexamethasone
AZD2906
Hydrocortisone
Mapracorat
AZD9567
GW870086
ZK216348
CORT108297
RU486
CpdA
B
num_dREs
max_LFC
mean_LFC
median_LFC
1
0.79
0.73
0.78
0.79
1
0.83
0.82
0.73
0.83
1
0.99
0.78
0.82
0.99
1
Pearson correlation
1
0.9
0.8
0.7
C
Correlation with PCs (STARR−seq)
treatment
replicate
molecule
−0.97***
−0.15
0.11
0.19
0.39**
0.9***
PC1
PC2
0.8
0.4
0.0
−0.4
−0.8
D
Wilcoxon, p = 0.0039
RMSE
0.3
0.2
0.1
0.0
RNA
STARR
AZD2906
Hydrocortisone
Mapracorat
AZD9567
GW870086
ZK216348
CORT108297
RU486
CpdA

Figure S3. Performance of regulatory elements with differential activity for regression and principal component analysis, related to Figure 3. A. Distribution of STARR-seq effect sizes for all 49,711 regions with significant activity. B. Correlation between regulatory efficacy and effect sizes for all regulatory elements with differential activity (dREs). C. Correlations between principal components and experimental variables. D. Comparison of residual mean squared error (RMSE) for the linear regression models in Figure 2 and Figure 3.

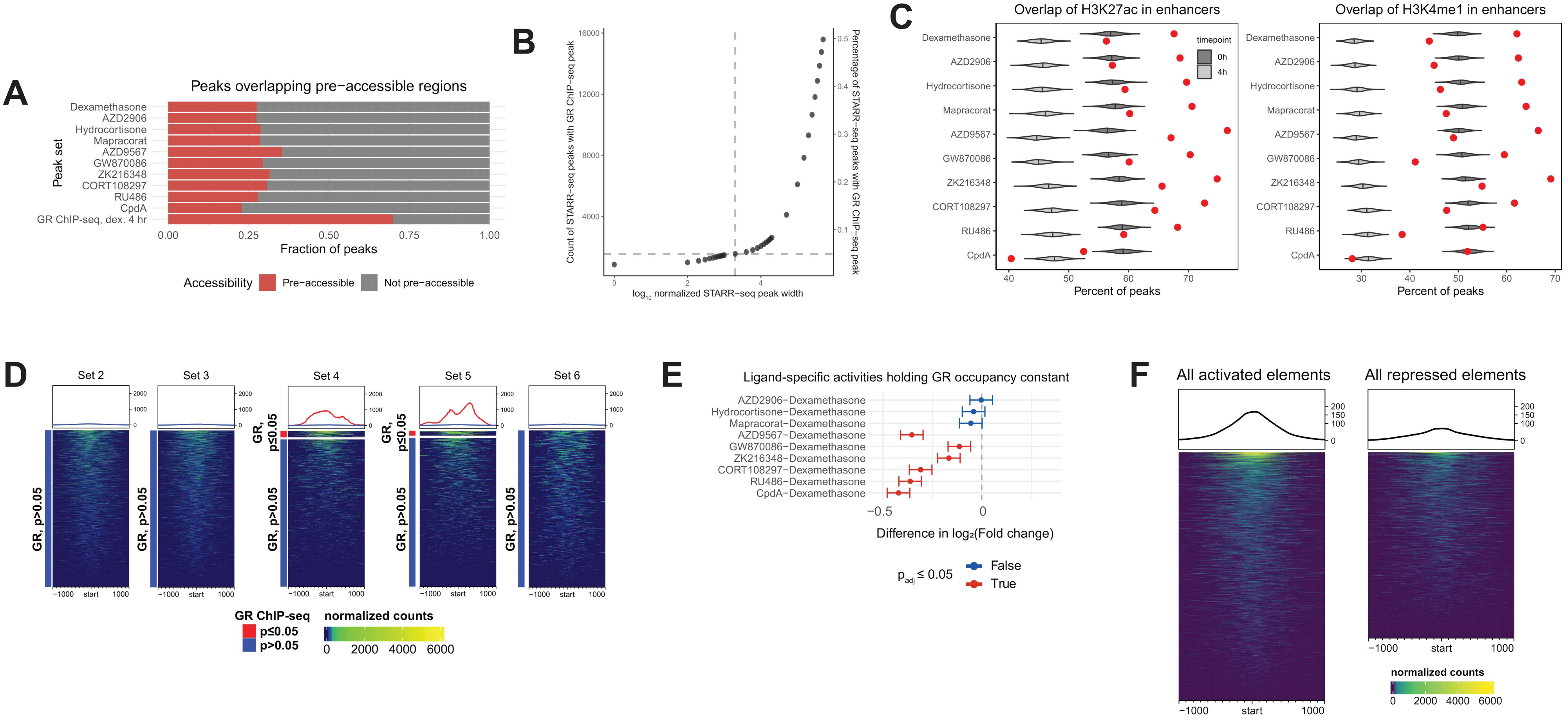

A
Peaks overlapping pre-accessible regions
Dexamethasone
AZD2906
Hydrocortisone
Mapracorat
AZD9567
GW870086
ZK216348
CORT108297
RU486
CpdA
GR ChIP-seq, dex. 4 hr
Peak set
Fraction of peaks
Accessibility
Pre-accessible
Not pre-accessible
B
Count of STARR-seq peaks with GR ChIP-seq peak
Percentage of STARR-seq peaks with GR ChIP-seq peak
log10 normalized STARR-seq peak width
C
Overlap of H3K27ac in enhancers
Overlap of H3K4me1 in enhancers
timepoint
0h
4h
Percent of peaks
D
Set 2
Set 3
Set 4
Set 5
Set 6
GR, p>0.05
GR, p≤0.05
GR ChIP-seq
p≤0.05
p>0.05
normalized counts
E
Ligand-specific activities holding GR occupancy constant
AZD2906−Dexamethasone
Hydrocortisone−Dexamethasone
Mapracorat−Dexamethasone
AZD9567−Dexamethasone
GW870086−Dexamethasone
ZK216348−Dexamethasone
CORT108297−Dexamethasone
RU486−Dexamethasone
CpdA−Dexamethasone
Difference in log2(Fold change)
padj ≤ 0.05
False
True
F
All activated elements
All repressed elements
normalized counts

Figure S4. Chromatin context for STARR-seq peaks with regulatory activity, related to Figure 4. A. Proportion of STARR-seq peaks and GR ChIP-seq peaks that overlap regions of pre-accessible chromatin (e.g., ATAC-seq peaks in the absence of GR ligand). B. Proportion of STARR-seq peaks with normalized length that overlap a GR occupied site (e.g., ChIP-seq peaks following 4 hr. dexamethasone treatment at 100 nM). C. Percentage of accessible STARR-seq peaks that overlap covalent histone modifications (e.g., ChIP-seq peaks following 4 hr. dexamethasone treatment at 100 nM). The distribution of percentage overlap between background regions and histone modifications is shown as a violin plot for each timepoint. The percentage of overlap between ligand-responsive elements and histone modifications is indicated with a red circle. Z-scores were calculated based on these measurements. D. Detected GR occupancy overlapping regulatory elements with different intersections of chromatin markers from Figure 4. E. Confidence internals estimated for differential ligand-responsive activity in Set 1 regulatory elements using the Tukey HSD test. Intervals that span the origin indicate no significant difference in activity due to ligand treatment. F. GR occupancy signal measured by ChIP-seq for all activated (left) and repressed (right) regulatory element loci.

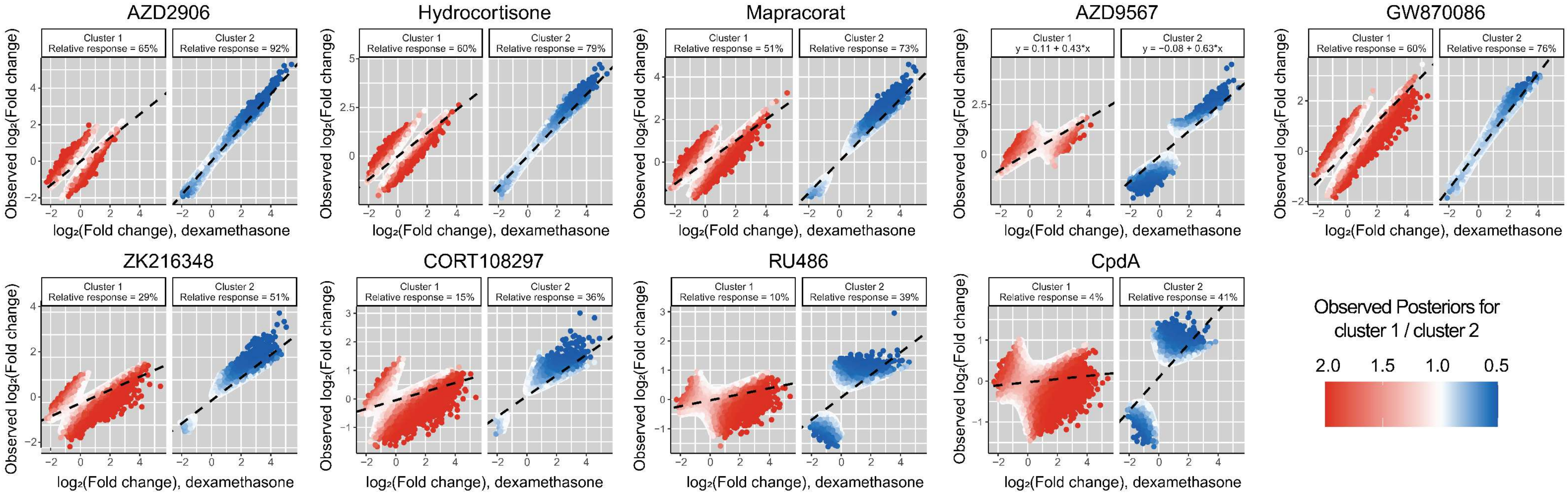
AZD2906
Cluster 1 Relative response = 65%
Cluster 2 Relative response = 92%
Hydrocortisone
Cluster 1 Relative response = 60%
Cluster 2 Relative response = 79%
Mapracorat
Cluster 1 Relative response = 51%
Cluster 2 Relative response = 73%
AZD9567
Cluster 1 y = 0.11 + 0.43*x
Cluster 2 y = −0.08 + 0.63*x
GW870086
Cluster 1 Relative response = 60%
Cluster 2 Relative response = 76%
ZK216348
Cluster 1 Relative response = 29%
Cluster 2 Relative response = 51%
CORT108297
Cluster 1 Relative response = 15%
Cluster 2 Relative response = 36%
RU486
Cluster 1 Relative response = 10%
Cluster 2 Relative response = 39%
CpdA
Cluster 1 Relative response = 4%
Cluster 2 Relative response = 41%
Observed log2(Fold change)
log2(Fold change), dexamethasone
Observed Posteriors for cluster 1 / cluster 2
2.0 1.5 1.0 0.5

Figure S5. Observed regulatory element responses using a two-component Gaussian mixture model for clustering, related to Figure 5. Elements are assigned to each cluster according to the maximum posterior probability, and relative responses for each cluster are estimated using linear regression.
